# Parallel clonal and molecular profiling of hematopoietic stem cells using RNA barcoding

**DOI:** 10.1101/2022.05.16.491933

**Authors:** Edyta E. Wojtowicz, Jayna Mistry, Vladimir Uzun, Anita Scoones, Desmond W. Chin, Laura Kettyle, Francesca Grasso, Allegra M. Lord, Graham Etherington, Charlotte Hellmich, Petter S. Woll, Mirjam E. Belderbos, Kristian M. Bowles, Claus Nerlov, Wilfried Haerty, Leonid V. Bystrykh, Sten Eirik W. Jacobsen, Stuart A. Rushworth, Iain C. Macaulay

## Abstract

Anucleate cells - platelets and erythrocytes - constitute over 95% of all hematopoietic stem cell (HSC) output, but the clonal dynamics of HSC contribution to these lineages remains largely unexplored. Here, we use lentiviral RNA cellular barcoding and transplantation of HSCs, combined with single-cell RNA-seq, for quantitative analysis of clonal behavior with a multi-lineage readout - for the first time including anucleate and nucleate lineages. We demonstrate that most HSCs steadily contribute to hematopoiesis, but acute platelet depletion shifts the output of multipotent HSCs to the exclusive production of platelets, with the additional emergence of new myeloid-biased clones. Our approach therefore enables comprehensive profiling of multi-lineage output and transcriptional heterogeneity of individual HSCs, giving insight into clonal dynamics in both steady state and under physiological stress.

## Main Text

Hematopoietic stem cells (HSCs) are rare bone marrow (BM) resident multipotent cells with potential to self-renew and to replenish all nucleated [1] and anucleate blood cells [2], sustaining long-term multilineage reconstitution after physiological and clinical challenges including chemotherapy and bone marrow transplantations [3,4]. Allogeneic HSC transplantation remains the only curative treatment modality for numerous hematological malignancies, however inefficient blood lineage replenishment, especially neutropenia and thrombocytopenia remain a major cause of morbidity and mortality [5,6]. Single cell transplantation experiments have uncovered significant heterogeneity among reconstituting HSCs, which may reflect different propensities for lineage commitment by distinct myeloid-, lymphoid-and platelet-biased HSCs [2,7–10]. Given such a high level of heterogeneity, it is unclear how different HSCs coexist in a polyclonal environment and how many HSC clones are simultaneously actively supporting production of nucleated and anucleate cells. Furthermore, although there are suggestions that platelet-biased HSCs may represent a fast-track to platelet production, the response of individual long term HSCs (LT-HSCs) and their clonal progeny to acute depletion of the platelet lineage remains unexplored. Furthermore, the transcriptional landscape underlying these clonal changes is understudied.

Quantitative clonal analysis depends on the method of labelling and detection. Cellular barcoding method was recently implemented for the clonal analysis of LT-HSC using genomic DNA [11–13]. The approach provided detailed information about LT-HSC subsets and clonal kinetics during haematopoiesis after transplantation [12,14], upon a challenge [15] or in aged animals [16]. Recently RNA-based barcoding has been applied for studies of the nucleated cells in the BM, however the labelling protocol applied did not effectively label platelets and erythrocytes [17], and thus a major component of HSC output was overlooked.

Here, for the first time, we have successfully barcoded both nucleate and anucleate lineages (including platelets) for kinetic clonal studies in PB. We applied RNA cellular barcoding method to FACS-purified LT-HSCs in combination with single-cell RNA sequencing (scRNA-seq) to perform kinetic clonal tracking of nucleated and anucleated blood cell populations in the hematopoietic system and linked it with molecular signatures of LT-HSCs, platelet and erythroid progenitors. We have taken advantage of the lentiviral library, where the barcode is located in the 3’ end of eGFP transcript, and is therefore present in both the genome of the cells and expressed as mRNA [18]. The approach is versatile, sensitive and applicable to other model systems including human cord blood or bone marrow cells [18].

Through cellular barcoding and transplantation, we have characterized clone size, developmental potential, and homing ability in almost 600 LT-HSCs. Our data provide detailed insight into functional differences and the dynamics of LT-HSCs response to stress. We demonstrate that most transplanted clones consistently contribute to hematopoiesis and have multipotent potential to produce myeloid, lymphoid, erythroid and platelet progeny, supporting the clonal stability model of hematopoiesis. Importantly, upon acute platelet depletion we observe the most striking changes in the myeloid lineage due to reprogramming of multipotent clones, which start to exclusively produce platelets and the emergence of new clones producing only myeloid progeny to compensate for the myeloid output of reprogrammed, multipotent clones. Our single cell RNA-seq analysis revealed that genes differentially expressed between reprogrammed and newly emerged clones are involved in the ribosome biogenesis, oxidative phosphorylation and chromatin remodelling.

## Results

### RNA barcoding combined with high-throughput sequencing enables quantitative clonal analysis

Before *in vivo* analysis of clonal composition in anucleate and nucleated cells we validated the sensitivity and accuracy of RNA and DNA barcoding assay *in vitro* using the K562 cell line as a model. DNA served as a positive control, being a widely accepted approach for *in vivo* lineage tracing [13,14,19]. Thus, defined mixtures of clonally expanded barcoded cells with previously published library [18] were analyzed in several dilutions ranging from 1 up to 5000 cells per barcode (Figure 1A). Using G&T seq we simultaneously isolated RNA and DNA from the same sample [20]. Individual samples were amplified with primers containing a unique multiplex tag and pooled multiplexed PCR products were sequenced (Figure 1A). The sequencing results were analyzed using previously published protocol [21], Materials and Methods). When we correlated read counts for the barcodes in cDNA and gDNA samples we observed high correlation between barcode abundance at different dilutions (R^2^=0.993) using generalised linear model fit (Figure 1B). We reliably detected barcodes in as few as 1 cell (Figure 1) and clones that represent only 0.01% of the starting population were successfully detected, while they were missed at DNA level (Figure 1B). Relative measurement error of barcode detection at DNA level was 13%, while for cDNA was 16%. Thus, cDNA provides superior sensitivity compared to DNA, however the barcode detection error is slightly higher, possibly related to the increased frequency of errors being introduced by the reverse transcriptase during the cDNA synthesis [22].

**Figure 1.**
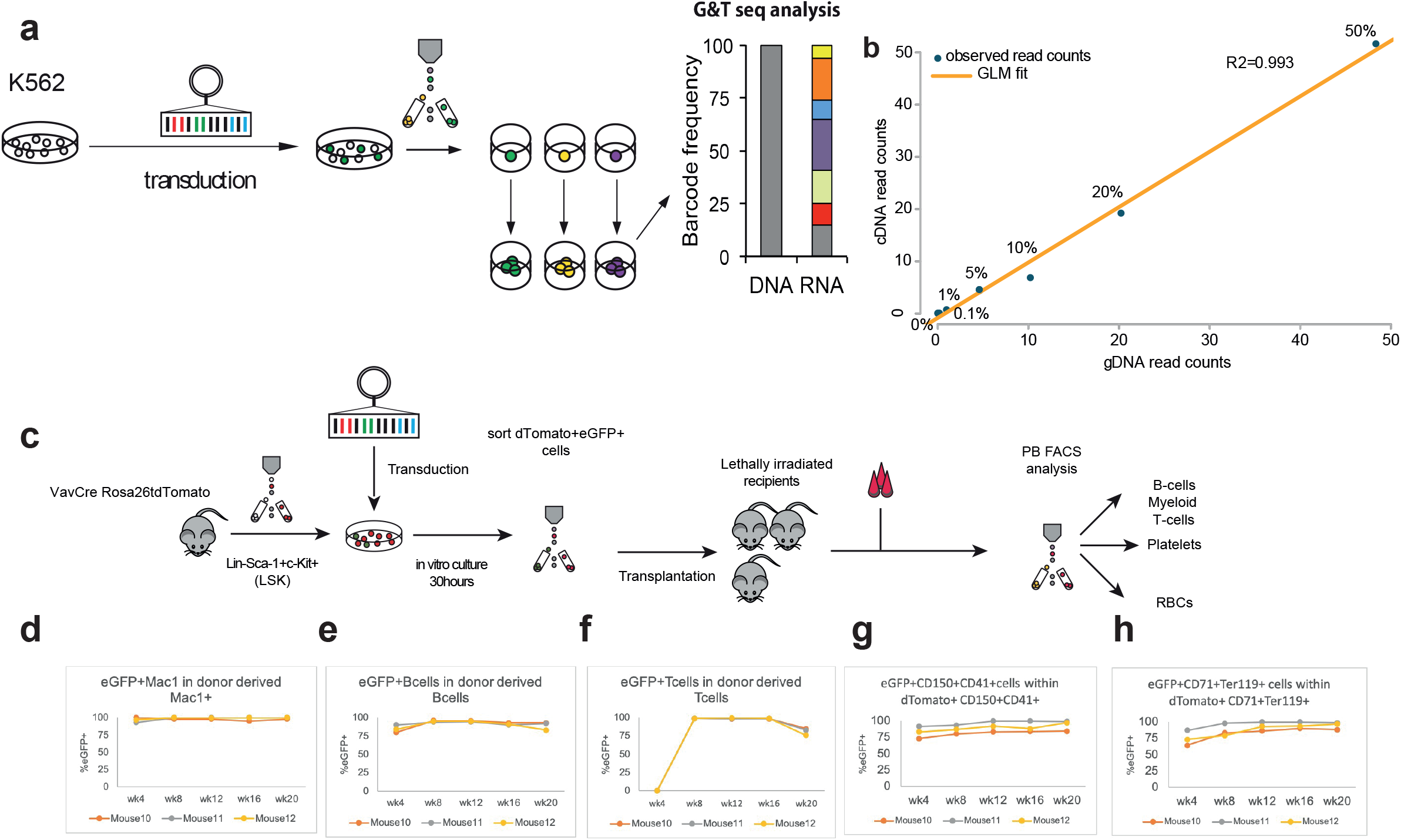
In vitro and in vivo validation of RNA barcoding approach for stable, long-term clonal kinetics studies of hematopoiesis. A.Experimental set up for *in vitro* validation of quantitative RNA based clonal studies. The K562 cell line was used for the transduction, single eGFP+ cells were FACS-sorted and expanded to generate monoclonal calibration samples. Four clones were used to create samples, where cells carrying different barcodes were mixed in known ratios-spiked in clone contributing 0.1%, 1%, 5%, 10%, 20% or 50% of total mix. B. Generalised linear model (GLM) correlation between the read count for barcoded transcript retrieved in calibration samples from cDNA or gDNA. C. Experimental setup to test the silencing of eGFP in VavCre x Rosa26tdTomato LSK cells D.-H. eGFP+ chimerism changes in tdTomato+ anucleate cells and CD45.2+ (donor derived) nucleated cells. To calculate the silencing in specific lineage we subtracted eGFP^+^ cells from CD45.2^+^ cells (nucleated cells-CD19^+^, Mac1^+^ or CD4/8^+^cells) or from tdTomato^+^ cells (all donor derived anucleate cells), shown are results from a single experiment in 3 recipient mice

Having confirmed limited skewing of the barcode representation at the RNA level at transduction efficiency below 50% we pursued *in vivo* validation of the labelling efficiency in nucleated and anucleate cells. We applied the Vav x Cre Rosa26 tdTomato mouse model, where all hematopoietic cells including platelets are tdTomato+ [23]. In brief, FACS-purified lineage-Sca-1+ c-Kit+ cells (LSK), were lentivirally transduced and after 24-hours we FACS purified tdTomato+eGFP+ cells and transplanted them into lethally irradiated recipients (Figure 1C). We analyzed blood cell contribution of transplanted cells and observed chimerism level reaching 85-100% at week 16, which started slightly decreasing at week 20 in T-cells. Importantly, we did not observe a loss of labelling in any particular lineage with the average silencing at 7.5% in all analyzed mature blood lineages (Figure 1D-H), indicating the approach is appropriate for sensitive barcode detection in nucleated and anucleate LT-HSC progeny in long-term transplantation studies.

### Clonal composition of erythroid, myeloid cells, B cells and platelets is stable over time

To study the clonal kinetics in platelets and nucleated cells *in vivo*, we transplanted 2,500 transduced barcoded LSK CD48-CD150+ cells (Suppl. Fig. 1A) into 4 busulfan treated recipients and measured eGFP+ chimerism (Figure 2A). PB reconstitution was first assessed by measurement of eGFP+ chimerism in platelets, RBCs, myeloid and lymphoid cells at 12 and 20/28 weeks by FACS (Suppl. Fig. 1 B-E).

**Figure 2.**
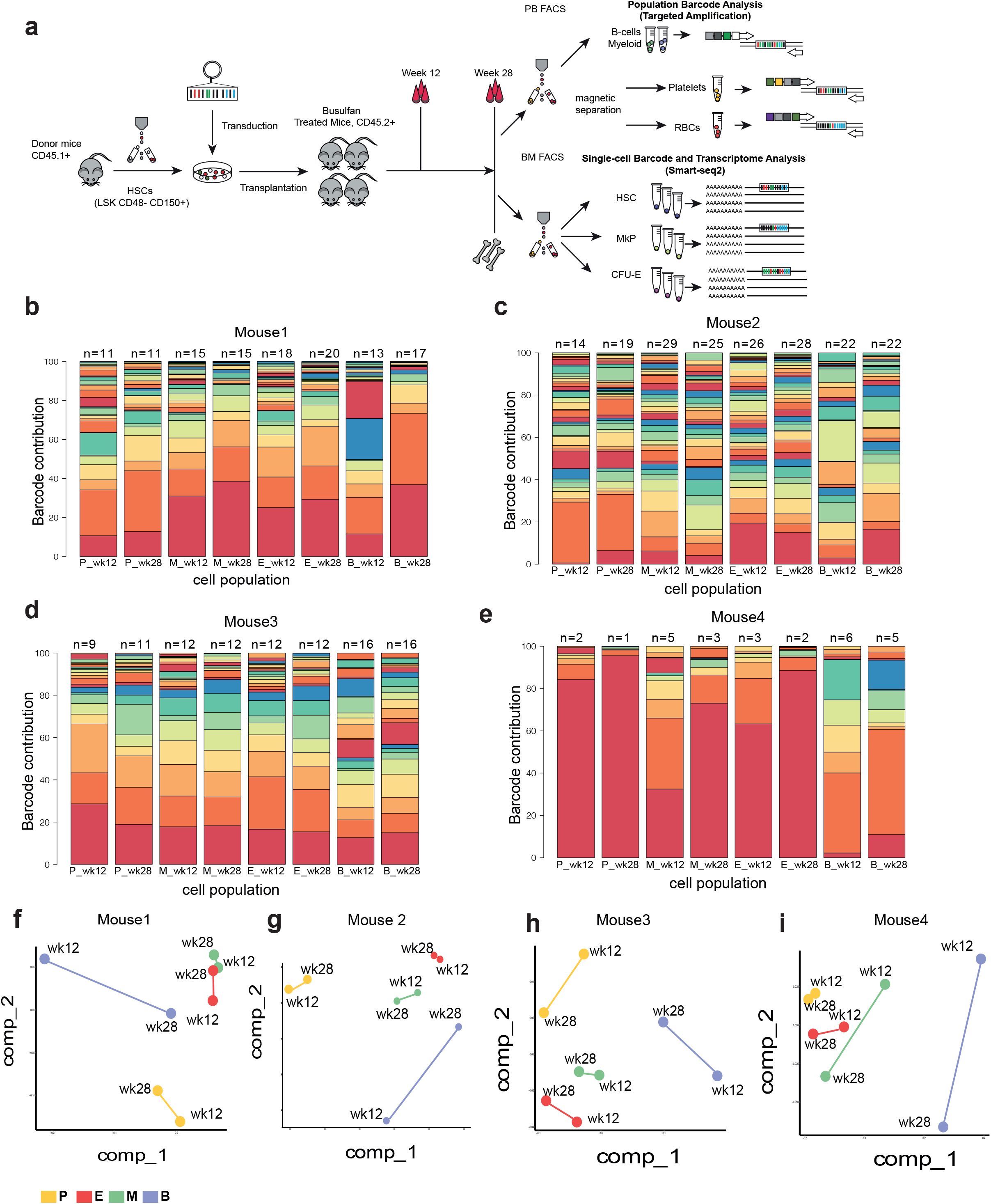
Clonal tracking in nucleate and anucleate blood cells. A. Experimental design for in vivo multilineage RNA barcoding and single-cell (BM) and bulk (PB) readouts (for 2 independent experiments). B.-E. Stacked bar plots representing clonal composition in PB populations (P-platelets, M-myeloid cells, E-erythroid cells, B-B cells) at 12 and 28 weeks in 4 analysed control animals, on top depicted is the Shannon count. F.-I. Multidimensional scaling (MDS) with Pearson correlation between barcode abundance (top 90%) in mature PB populations (P-platelets, E-erythroid cells, M-myeloid cells, B-B cells) at 12-and 28-weeks post transplantation.

Subsequently, RNA from FACS purified myeloid cells and B-cells (over 99% purity, Suppl. Fig. 2A-C, Table 1A), bead purified RBCs (purity 99.84±0.045%) and platelets (purity 99.96±0.023%) at weeks 12 and 20/28 post transplantation (Table 1B-C) was used as input for barcode quantification by sequencing. Replicate samples from each lineage of the same mouse displayed a high correlation of detected barcodes (mean R^2^ = 0.95, Suppl. Fig. 1F-I), indicating robust barcode detection in analysed samples. This was especially important for platelet samples, since RNA content of these is 10,000 times lower compared to nucleated cells [24], therefore this validation step reassured robust barcode capture also in platelets. To perform the clonal analysis in PB we ranked barcodes from most to the least abundant in blood populations in each mouse and identified the ‘dominant’ set of barcodes which collectively accounted for >90% of total abundance and focused our clonal kinetic analyses on these barcodes [21,25]. We were able to detect clones contributing to platelet lineage at 0.01_±_0.11, erythroid 0.03_±_0.05, myeloid 0.02_±_0.1, B cell 0.02_±_0.12 (Suppl. Figure 1 J-M, Table 2).

We used Shannon count to estimate a number of contributing clones in addition to a classic assessment of the clonal diversity within lineages estimated by Shannon diversity index [21]. We detected between 2-29 (week 12) and 1-28 (week 28) PB clones per mouse, with the number of clones dependent on the lineage analysed the lowest count in platelets and the highest in RBCs (Figure 2B-E). The stable contribution of individual clones to PB output (detected at both time points) was highly variable - we detected large clones which contributed 10.6%-72% of PB output, while the smallest detected multilineage clones contributed less than 1% to all lineages (Suppl. Table 2).

The kinetic changes of barcode composition in PB lineages and timepoints were summarised using multidimensional scaling (MDS) with Pearson correlation as a measure of similarity between barcode abundance in different PB lineages (Fig. 2F-I). These analyses demonstrate that clonal composition within cells of the same lineage is stable over time. In line with previous findings [2,16], fluctuations of clone size could be observed with some variation at week 12 and barcode composition becoming more similar among lineages at later time points (Fig. 1B, Suppl. Fig. 3A-D). This suggests that at early time points, we observe clones produced by shorter-living, transient progenitors supporting PB lineages. This is especially evident in B cells due to their lower turnover rate when compared to platelets, myeloid cells or even erythroid cells, but the emerging similarity at later time points indicates that LT-HSCs are then supporting all PB lineages including long-living B-cells. Thus, our approach enables sensitive and stable detection of clonal output in nucleate and anucleate PB lineages. At the same time, tracking clones for longer time periods was essential for understanding the dynamics of clonal fluctuations.

### Multipotent clones are the most abundant and productive in the bone marrow during steady state hematopoiesis

To further investigate the relationships between the clonal composition in PB and BM progenitor populations, we performed single-cell RNA-seq using Smart-seq2 [26] on eGFP+ BM HSC and progenitors giving rise to anucleate lineages (LT-HSCs, MkPs and CFU-Es) at 28 or 32-weeks post transplantation (Figure 1A, Suppl. Figure 3A). This enabled comprehensive, full-length transcriptomic profiling of 2,569 single cells from these phenotypically defined populations with parallel recovery of expressed cell barcodes in over 90% of cells (Fig. 3A, Table 4). Each animal contained between 2-49 uniquely barcoded LT-HSC clones.

**Figure 3.**
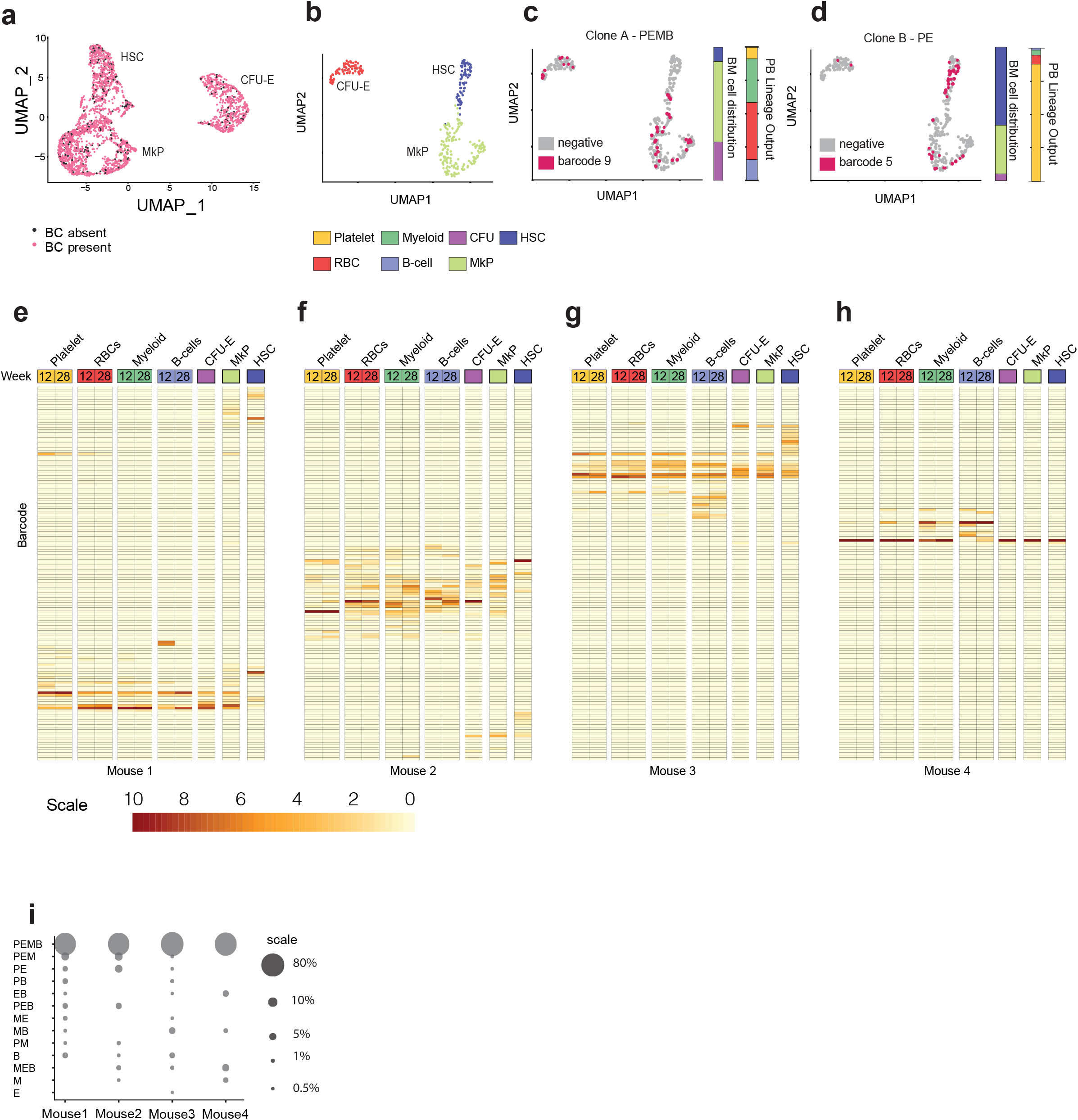
Clonal tracking in blood and bone marrow cells exerts high overlap between detected clones. A. UMAP plot showing single cell transcriptomes of sorted BM populations in which barcodes (BC) were detected (all cells which had over 50 000 reads and mitochondrial gene count<10% with barcode read count > 3 were included) B. UMAP plot showing cell clusters based on single cell transcriptomes of BM populations in mouse 2 (32-weeks post transplantation). C. UMAP plot representing clone A-cells carrying barcode rank 9 in mouse 2 in the BM, bar plot representing PB lineage output (multilineage - PEMB, contribution to all 4 lineages >0.1% at 2 consecutive time points) and the distribution of this barcode within the BM cell types profiled. D. as for C, but clone B is shown. This clone was classified as having more platelet-erythroid restricted output (PE, contribution to P and E lineages>0.1% at 2 time points). E -H. Heatmaps showing the Spearman correlation between abundance of dominant unique barcodes (rows) in PB (Platelets, RBCs-erythroid cells, Myeloid, B-cells) and BM populations (HSC, MkP, CFU-E) clones in 4 control animals (separated columns) at week 12 and 28 (PB) and 32 (BM) post transplantation. I.Bubble plot depicting the cumulative PB contribution of different types of clones in 4 control animals (Mouse 1-4). For the analysis only dominant barcodes were included. To consider the clone as contributing to the lineage it had to contribute >0.1% at 2 time points. Abbreviations: PEMB: platelet-erythroid-myeloid-B-cell, PEM: platelet-erythroid-myeloid, PE: platelet-erythroid, PB: platelet-B-cell, EB: erythroid-B-cell, PEB: platelet-erythroid-B-cell, ME: myeloid-erythroid, MB: myeloid-B-cell, PM: platelet-myeloid, B: B-cell, MEB: myeloid-erythroid-B-cell, M: myeloid, E: erythroid.

We subsequently mapped PB lineage output onto single FACS-purified and sequenced HSCs, MkPs and CFU-E transcriptomes (Fig. 3B-D, Suppl. Figure 4A-D). This allowed classification of individual HSCs based on the composition of their lineage output (stable contribution > 0.01% to the lineage at two time points), as shown here for two clones from mouse 2 - clone A which demonstrated multilineage Platelet/Erythroid/Myeloid/B-cell (PEMB) output and clone B which showed more restricted Platelet/Erythroid (PE) output. To classify the clone as eg. PEMB it had to contribute >0.01% to all 4 lineages at 2 time points, PE had to contribute >0.01% to P and E lineages, but not other lineages at 2 time points.The overlap between PB and BM clones revealed up to 70% of shared clones, however they had different contributions to blood production and abundance in the BM (Figure 3 E-H).

The analysis of barcode overlap between CFU-E and blood erythroid cells or MkP and circulating platelets revealed that 45.2 ± 12.1% of erythroid and 40.55 ± 15.7% of platelet clones would have been missed if the analysis would include only clones present in nucleated progenitor cell populations in BM (Figure 3 E-H, Suppl. Figure 3B-E). This disparity may relate to limited sampling of the *in vivo* progenitor cell populations, although it is possible that some of these cells may be generated via a bypass of the MkP stage [27]. Similarly, MkP/CFU-E progenitors may become exhausted, but leave progeny that is still detectable in PB. Therefore, the most reliable clonal kinetics study of PB is a direct analysis of nucleated and anucleate cells present in the circulation.

Efficient barcode recovery from BM and PB lineages enabled parallel analysis of clonal composition in nucleated and anucleate cell types and their progenitor cells in individual mice (Fig. 3 E-H). In each mouse, a fraction of clones found in the PB lineages could be found in the BM stem/progenitor cell populations [2]. We have observed functional HSC heterogeneity in their reconstitution patterns. In mice 1-3, 40% of PB clones overlap with barcodes detected in progenitors, and the overlap is slightly lower with HSCs (30%, Fig. 3E-H). We also detected an extreme case of oligoclonality in mouse 4, where 70% of the PB and over 80% of the BM cells shared the same barcode (Figure 3H). Remaining clones present in PB, not detected in BM may reflect longevity of other cell types supporting PB production at the time of our analysis or might result from limited BM sampling in our study (potentially lower abundance or asymmetrical distribution of these clones in the BM). Overall this data demonstrates the stochasticity in clonal composition of PB and BM cell populations in different animals transplanted in the same experiment.

Next, we have summarized the clonal output of PB clones during steady state hematopoiesis (Figure 3I) and observed the highest output to mature blood lineages had multipotent clones.

Clones only supporting erythroid cells and B-cells (EB) are likely reflecting the output from short-term HSCs or multipotent progenitors, since they were unable to sustain multilineage repopulation in the long-term (Suppl. Table 4B). Overall, multipotent PEMB clones were the most productive supporting almost 80% of all PB output (Fig. 1K) in agreement with previous findings in nucleated cells [11,28]. On the contrary platelet biased clones were significantly less abundant in PB and their contribution varied from 0.1% up to 10% at steady state following transplantation.

Our approach allowed recovery of over 90% of clonal information from single BM cells, with the average 1M reads/cell. Analysis of barcode composition in PB and BM populations revealed up to 70% of shared clones between both tissues, which is significantly higher compared to previous barcoding studies applying DNA [29]. The clonal composition among animals transplanted within the same experiment highlighted high heterogeneity and repopulating potential of LT-HSCs. Importantly, our analysis has stressed the disparity between the BM and PB clonal composition in anucleate cells and their BM progenitors, highlighting the need for the direct clonal assessment in these blood lineages, rather than using BM progenitors as a good approximation of clonal diversity in the blood.

### Platelet depletion forces repurposing of the lineage output in multipotent clones

Next, we wanted to evaluate how the hematopoietic system would respond to acute, but transient, depletion of platelets. To assess cellular and multilineage clonal changes in response to acute platelet depletion, we induced acute thrombocytopenia in mice (Figure 4A) transplanted with barcoded HSCs 28-weeks post transplantation (Suppl. Figure 5 A-D) [30]. We have confirmed the efficient platelet depletion 24-hours post treatment (Suppl. Figure 5E). The kinetics of platelet replenishment has been extensively studied and requires 6-10 days for platelets to reach normal counts [10].

**Figure 4.**
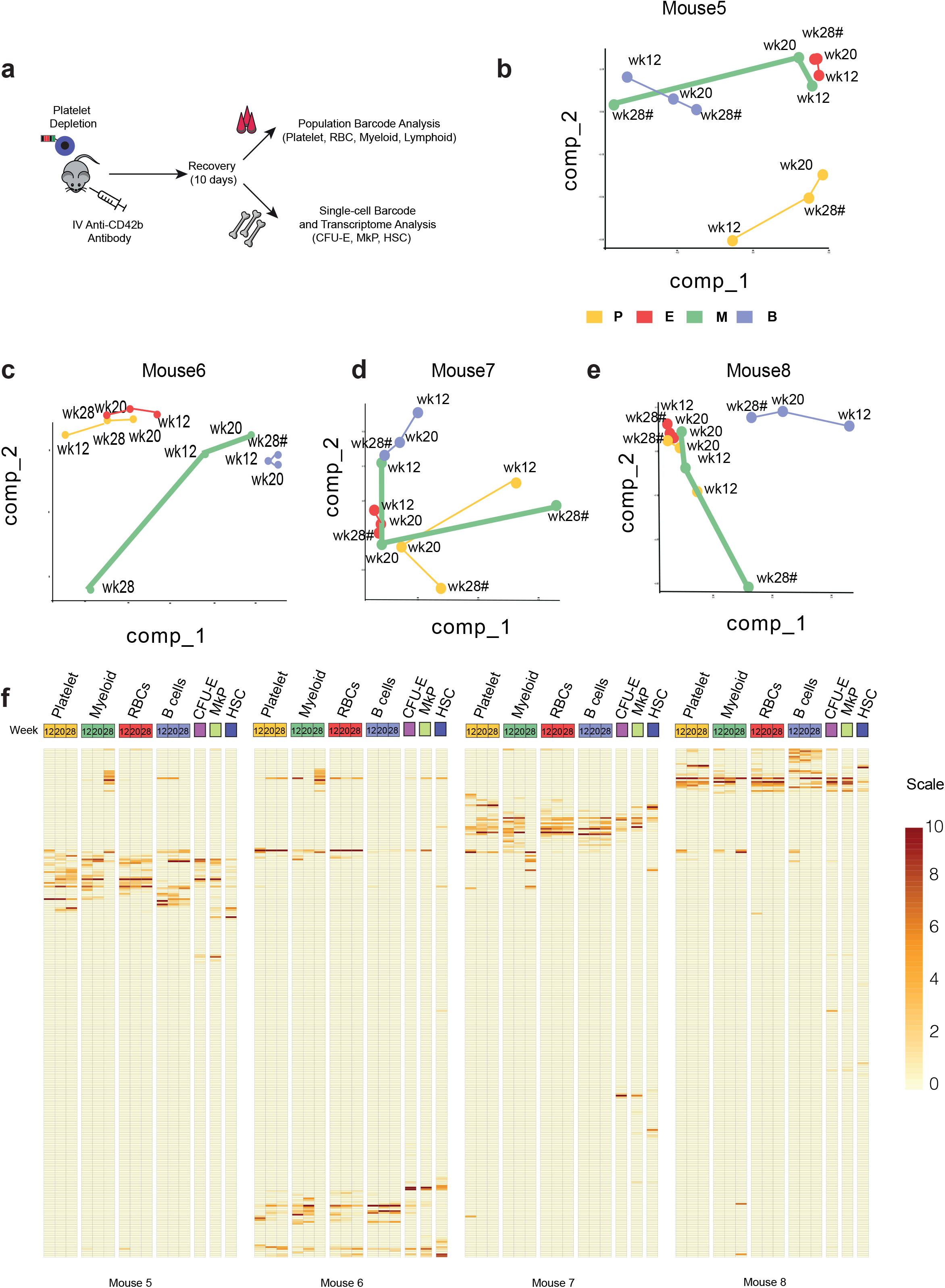
Acute platelet depletion causes dramatic changes in clonal composition of the myeloid lineage causing the recruitment of completely new clones. A. Experimental design for platelet depletion studies B. - E. MDS with Pearson correlation between barcode abundance in PB cell populations in (P-platelets, E-erythroid cells, M-myeloid cells, B-B-cells) at 12-, 20- and 28-weeks+10days post transplantation in Mouse 6 pre-(12, 20 weeks) and post-(28 weeks+10days) platelet depletion F. -I. Heatmaps showing the Spearman correlation between abundance of dominant unique barcodes (rows) in PB (Platelets, RBCs-erythroid cells, Myeloid, B-cells) and BM populations (HSC, MkP, CFU-E) clones in 4 platelet depleted animals (separated columns) at week 12, 20 and 28 (PB) and 28 (BM) post transplantation.

We collected BM and PB samples 10 days after treatment (28 weeks +10 days), at which point thrombocytopenia had resolved (Suppl. Figure 5E) and determined PB and BM progenitor clonality at bulk and single-cell resolution, respectively (Figure 4B-F). When we compared the similarity among PB lineages before and after the platelet depletion at two time points before (week 12 and 20) and one after the platelet depletion (week 28+10days) using MDS plots we observed striking changes in the clonal composition of the myeloid lineages, rather than the platelet lineages, as we would have expected (Figure 4B-E). Almost 90% of all multipotent clones contributing to myeloid cells at week 20 have stopped producing them and redirected their output to replenish platelets, which constituted 30-50% of newly produced platelets. This group of barcodes was also found in erythroid cells and B cells (Figure 4G, platelet-erythroid-Bcell clone, PEB clones), which can be explained by the half life time of the latter two cell types (22 days and 13-22 weeks respectively [31,32], strongly suggesting they have been produced before the platelet depletion, the time when the clone was multipotent and supported all analyzed blood lineages (Figure 4F, G).

Interestingly, we did not notice decreased count or frequency of myeloid cells 10 days after the platelet depletion. Myeloid cells have short half-life time in peripheral blood (12.5 hours to 7 days [33]), therefore require a constant input from upstream progenitor cells and HSCs. This suggests that new clones have been recruited to solely produce myeloid cells. Indeed, when we looked into the clonal composition of the myeloid cells the missing output from multipotent clones has been matched by a group of newly emerged clones exclusively producing almost 80% of myeloid cells in PB (Figure 4F).

### Molecular signatures of repurposed multipotent and newly emerged myeloid clones in LT-HSC

Since we have noticed striking clonal changes upon platelet depletion, we have taken the advantage of the scRNA seq-data from the BM populations to link single-cell transcriptional profiles with PB output. Firstly, we have performed unsupervised clustering of treated and non-treated LT-HSC, MkP and CFU-E. Cells from each type clustered together, regardless of experimental group, indicating that the changes in PB clonality is not reflected by a global shift in progenitor gene expression (Figure 5A, Suppl. Figure 5F). Next, we have performed supervised clustering based on the output of PB clones, which we linked with the molecular signature in LT-HSCs carrying the same barcode in the BM. We have compared differentially expressed genes between multipotent clones in control animals with P/PE/PEB clones in platelet depleted animals. This analysis revealed enrichment in GO terms such as ribosomal biogenesis, oxidative phosphorylation and platelet activation (*Gp9, Itga2b*). It suggests that LT-HSCs upon platelet depletion are translationally active and switch to more efficient oxidative phosphorylation to meet increased energetic demand during platelet replenishment, as suggested in a recent study [34]. Similar mechanisms of switched metabolism are observed in LT-HSCs during inflammation, when high platelet consumption is also observed[35]. Since the Itga2b (CD41) antibody was included in the purification strategy of MkPs, we reanalysed this data and observed increased protein expression of Itga2b in LT-HSCs from platelet depleted animals when compared to control ones (Figure 5C-D). This observation is consistent with a previous report showing an emergence of a new CD41+ HSC fraction after viral infection (pI:pC) during emergency thrombopoiesis [27,36].

**Figure 5.**
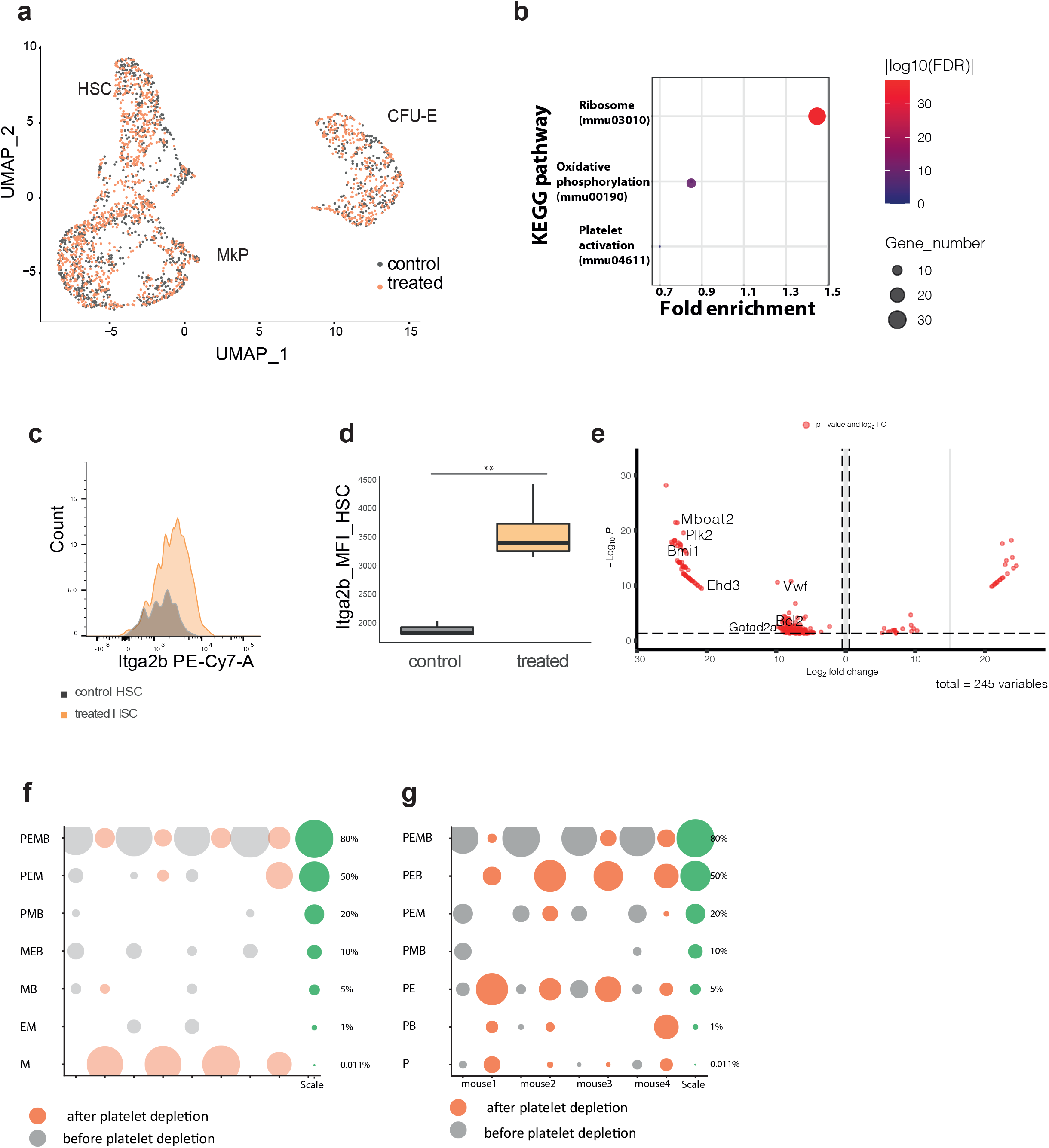
Depletion of platelets is also reflected in molecular signatures of HSCs with different blood output. A. UMAP plot representing the transcriptomic state of control and treated HSC, MkP and CFU-E (included cells with read count>50 000, mitochondrial gene content<10%) B. KEGG pathways enriched in platelet biased clones compared to control multipotent clones. C. Itga2b expression in LSK CD150^+^48^-^eGFP^+^ cells measured 28-weeks+10 days post transplantation in control and treated animals. D. Increased mean fluorescence intensity (MFI) of Itga2b in LSK CD150+48-eGFP+ in treated compared to control animals. E. Volcano plot representing genes upregulated in newly emerged myeloid clones compared to PEB clones F. Bubble plot representing the contribution of different clone types to the myeloid lineage before and after platelet depletion G. Bubble plot representing types and contribution of clones to the platelet lineage

Next, we asked the question what molecular changes are occurring between repurposed multipotent clones (PEB clones) and newly appeared myeloid clones. We filtered the scRNA-seq data set and selected only HSCs carrying barcodes identified in PB as PEB or new myeloid clones in the same animal. This analysis left 245 (p<0.001) differentially expressed genes. Newly emerged myeloid clones have increased expression of proteins involved in the chromatin remodeling (*Bmi1, Gatad2a, Mboat2*), cell cycle regulators (*Plk2*), antiapoptotic genes (*Bcl2*) and *Vwf* compared to PEB clones (Figure 5E). Of these, only five genes were differentially expressed between PEB and new myeloid clones in all animals: *Mboat2, Vwf, Ehd3, Bcl2* and *Gatad2a*. High expression of Vwf, a marker for the platelet lineage which is expressed in platelet-biased LT-HSCs, MkP, mature megakaryocytes and platelets [2,10] may suggest that these cells will start contributing also to the platelet lineage. Mboat2 is an acetyltransferase involved in the lipid metabolism, while Gatad2l is a member of the nucleosome remodeling and deacetylase complex (NuRD). Thus this signature may indicate these new myeloid clones are undergoing chromatin remodeling which underlies the changes in lineage output.

## Discussion

Detailed studies of HSC clonality in the hematopoietic system require reliable, stable and heritable marking of HSCs to trace their progeny in time. In this study, we have applied a novel RNA barcoding method to provide quantitative and dynamic tracking of individual HSCs with regard to their lineage contribution and transcriptional signature during steady state and upon acute platelet depletion.

In contrast to previous RNA barcoding studies of hematopoiesis [17], our approach labels both nucleated and anucleated PB lineages. We have validated RNA to be 100-fold more sensitive compared to a previously published DNA barcoding method [16] and quantitative source of clonal information, which should be a method of choice when studying hematopoiesis. We did not observe significant silencing of the barcode in labelled PB cells ensuring the robust, stable, long-term label retention in these cells allowing quantitative studies in not analyzed so far platelet lineage (Figure 1G). We have observed that almost 80% of PB output at steady state comes from multipotent clones, which produce 50-80% of all platelets in the PB. In our polyclonally repopulated recipients we observed significantly lower contribution of platelet biased clones at 0.5-4% of total platelets. It suggests that platelet biased LT-HSC are producing only a small fraction of all platelets in our model, which is different from single HSC transplantation studies [2]. Thus, it raises a possibility where platelet specific LT-HSC will only produce large quantities of platelets during extreme stress of the single stem cell transplantation, while when transplanted with multipotent LT-HSCs will contribute only a limited fraction of them.

Although multipotent clones were the most abundant in the BM, we have observed significant clonal diversity between the clonal composition in the BM progenitors (CFU-E and MkP) and their mature anucleated PB progeny, which reached 50%. This emphasizes the importance of including mature PB lineage analysis, rather than their BM precursors when studying the clonal kinetics in long-term PB studies. It also confirms that cell barcoding approaches capable of labelling all PB lineages are essential to observe the kinetics of stress responses in the haematopoietic system.

Acute platelet depletion revealed the remarkable plasticity and clonal dynamics of HSCs in response to the clinically relevant stress of thrombocytopenia. Here we were able to study how HSCs repurpose their blood output and observed a clear repurposing of multipotent clones prevalent at steady state to quickly and efficiently replenish platelets. It was accompanied by the emergence of completely new myeloid clones dominating upon platelet depletion. Based on literature one would expect that platelet biased HSC clones would quickly and significantly increase their PB output[2] being the major contributors, however our results show that it is easier for the system to repurpose the lineage output of 90% of already present multipotent clones towards platelets, empty the BM niche and recruit new group of HSCs to exclusively contribute to myeloid cells (Figure 6A). The need to replenish myeloid cells, but not B cells or erythroid cells is most likely related to their similar half-life time compared to platelets and likely existence of a shared upstream progenitor cell, which is unable to sustain increased platelet numbers and replenish myeloid cells at the same time.

**Figure 6.**
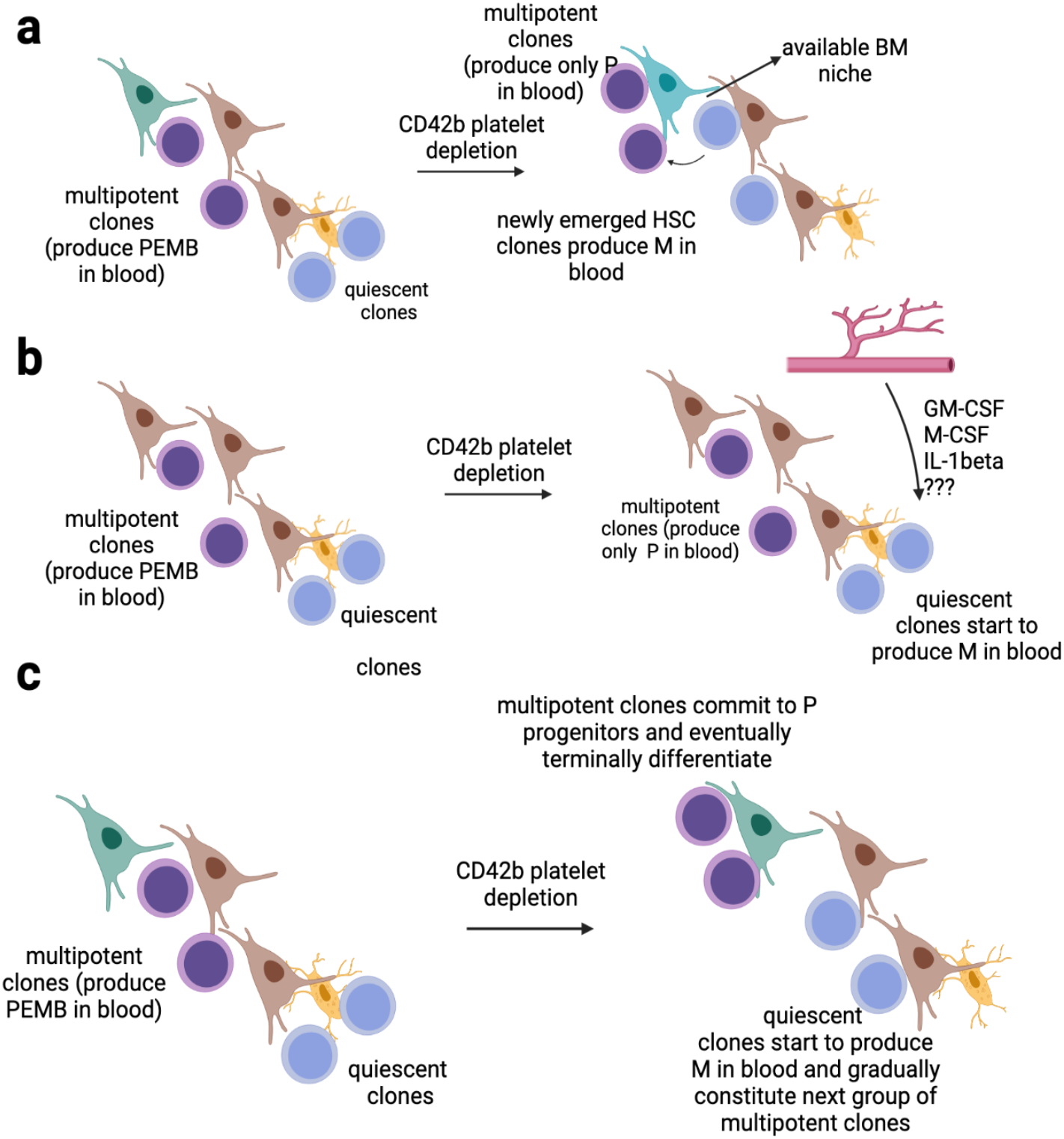
Three potential mechanisms for clonal changes in LT-HSCs following acute platelet depletion. A. At steady state, multipotent (PEMB) HSC clones occupy sinusoidal niches in the BM and produce nucleated and anucleated progeny in blood. Upon acute platelet depletion, multipotent clones shift their blood output, produce only platelets and shift the niche. Available BM niche is getting occupied by quiescent HSC clones, which did not support hematopoiesis before. These clones exclusively support the myeloid (M) lineage, however may start contributing to other lineages (eg. platelets and red blood cells) B. During steady state hematopoiesis multipotent clones (PEMB) produce all progeny in blood, upon platelet depletion they shift their output exclusively to replenish platelets. Platelets and myeloid cells share upstream progenitor cells, which are redirected to exclusively produce platelets. It causes short-term neutropenia, leading to the release of cytokines promoting myelopoiesis. These blood derived agents stimulate quiescent HSC clones in the bone marrow to contribute to the myeloid lineage and compensate for missing output from multipotent clones. C. Platelet depletion causes commitment of previously multipotent HSC clones (PEMB) to platelet progenitors, quickly and efficiently replenishing this lineage and emptying the BM niche. It causes quiescent HSC clones to enter the niche and start replenishing myeloid lineage (M). Gradually the first generation of multipotent clones (PEMB) differentiates and dies out and the second generation of multipotent clones starts to contribute also to the platelet lineage and eventually erythroid lineage.

High expression levels of *Vwf*, a marker of the platelet lineage, may suggest that these clones will start contributing also to platelets and potentially other lineages becoming the 2nd generation of multipotent clones, which so far has been only observed in the serial transplantation studies [17]. The difference between single HSC transplantation and our model may potentially be explained by the assay, where maximal repopulating potential of the HSC is being evaluated. On the contrary, in the polyclonally repopulated animals there is a pool of HSCs, which do interact with each other and coordinate their response upon stress. The latter scenario is closer to the actual situation in patients and therefore provides more physiological clues into HSC plasticity and response to stress.

In summary, our novel barcoding approach for the first time enables true multi-lineage tracking of hematopoietic output in both anucleate and nucleate lineages, in steady state and following a clinically relevant stress. Using scRNA-seq and barcode detection, effective in over 90% of single cells, the clonal output of individual cells can be linked with heterogeneity in gene expression, identifying molecular signatures of lineage biases. We demonstrate that acute platelet depletion enforces a switch in the clonal output from multipotent to exclusive platelet production, and the emergence of new, myeloid-biased clones.

## Supplementary Figures

**Supplementary Figure 1.**
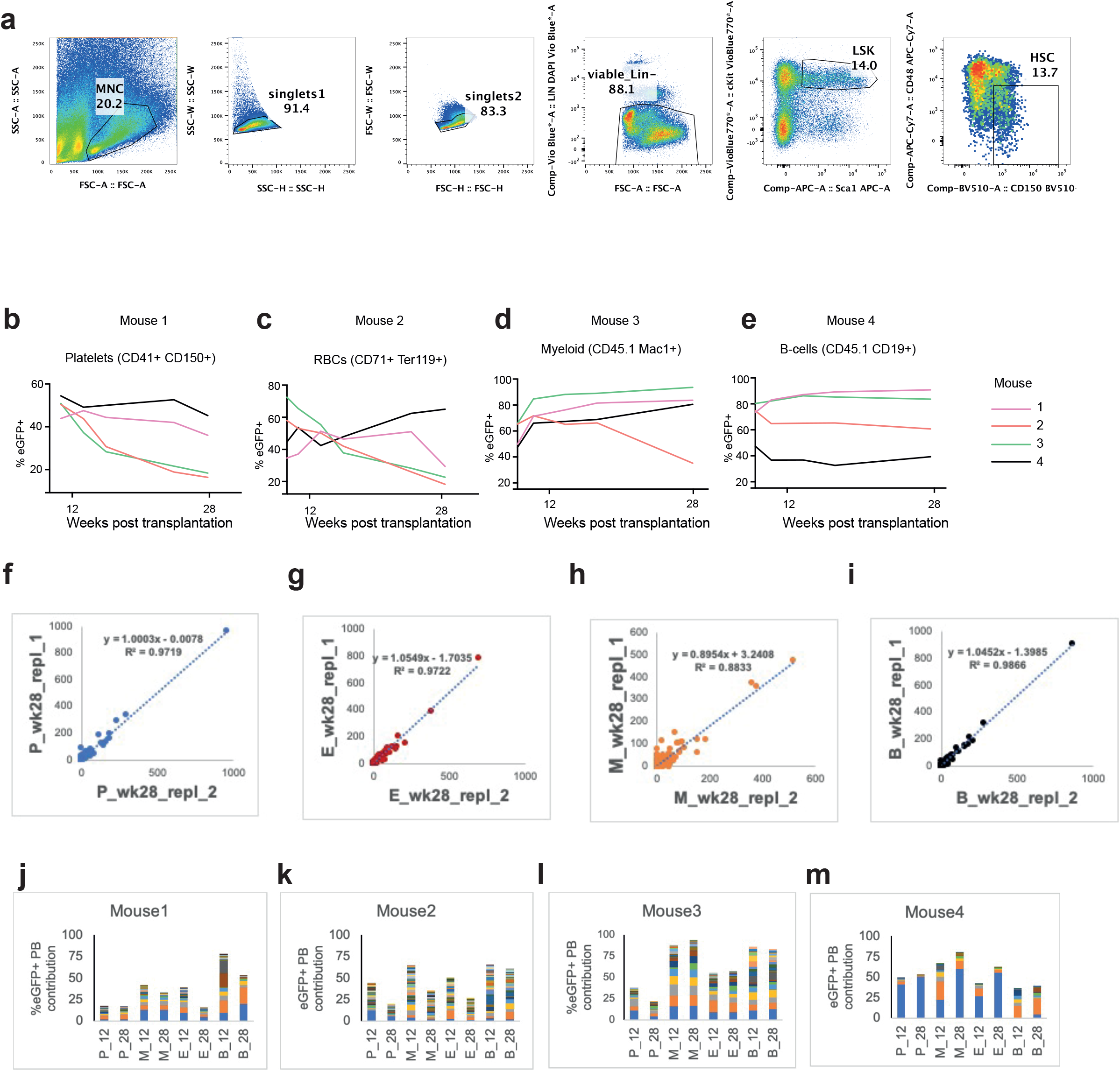
A. Gating strategy to purify HSC (LSK 48^-^CD150^+^) for transduction and transplantation experiments (4-6 mice used as donors in each experiment) B. - E. Contribution of transduced, eGFP^+^ HSC into 4 mature PB populations during the course of the experiment, plots are showing the chimerism level in mice 1-4 (corresponds to clonal composition in mice depicted in Fig.2 and 3) F. -I. Linear regression between 2 technical replicates for cDNA retrieved from 4 mature PB populations (P-platelets, E-erythroid cells, M-myeloid cells, B- B-cells) to assess the reproducibility of barcode recovery (displayed are normalised barcode read count in 2 technical replicates in the same animal) J.-M. Contribution of clones (top 90% of all barcodes detected in PB) corrected for the chimerism level in analysed PB lineages (corresponds to animals in Fig. 2 and 3)

**Supplementary Figure 2.**
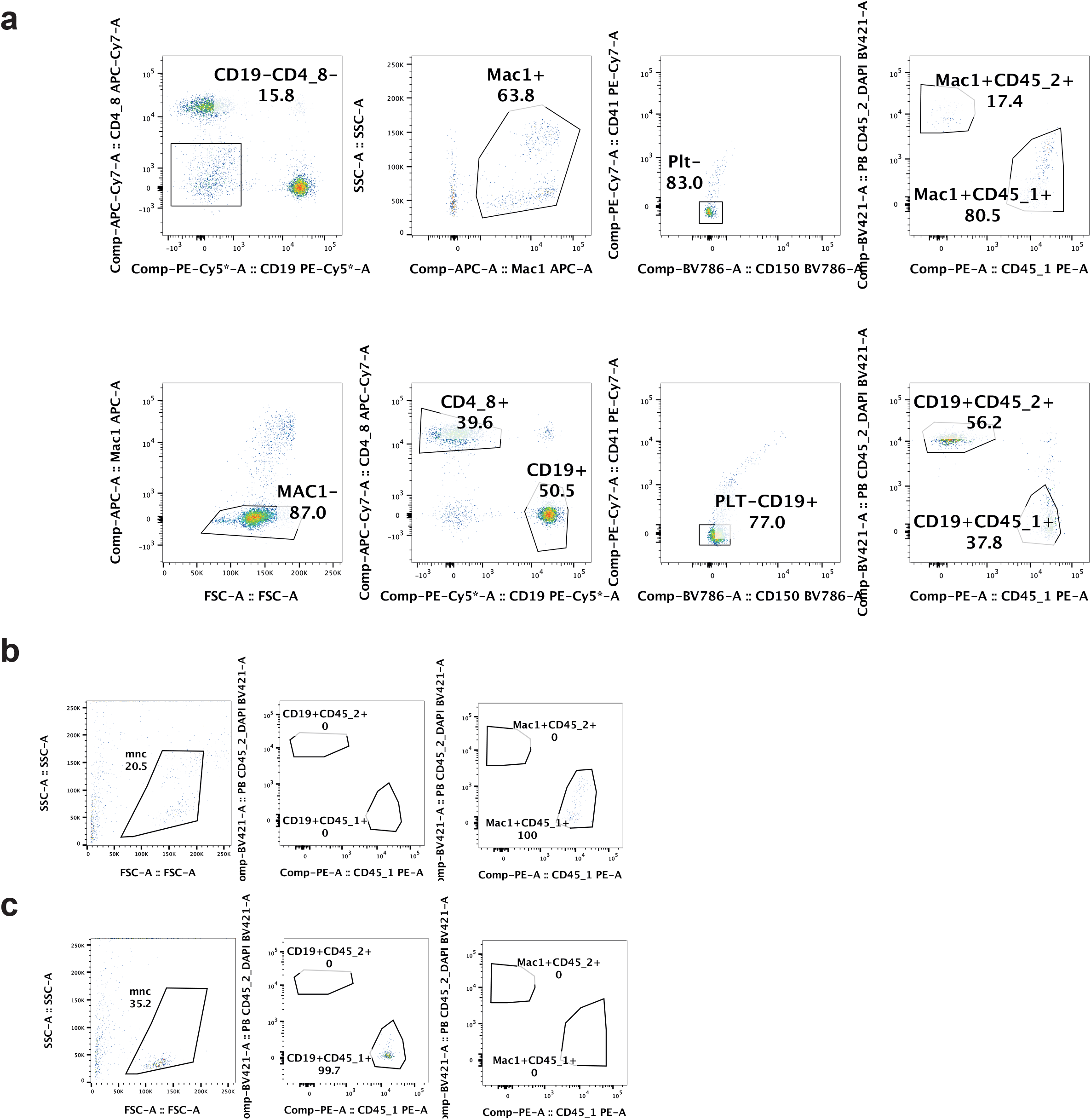
A. Gating strategy for PB cell population FACS-purification. Displayed are gates within the MNC/singlet/viable cell population. FACS definitions for purified populations: Mac1^+^ (CD19^-^CD4/8^-^CD150^-^CD41^-^CD45.1^+^), CD19^+^ (Mac1^-^CD4/8^-^CD19^+^CD150^-^CD41^-^CD45.1^+^) B. Purity of sorted Mac1^+^ cells evaluated by FACS Purity of CD19^+^ cells evaluated by FACS

**Supplementary Figure 3.**
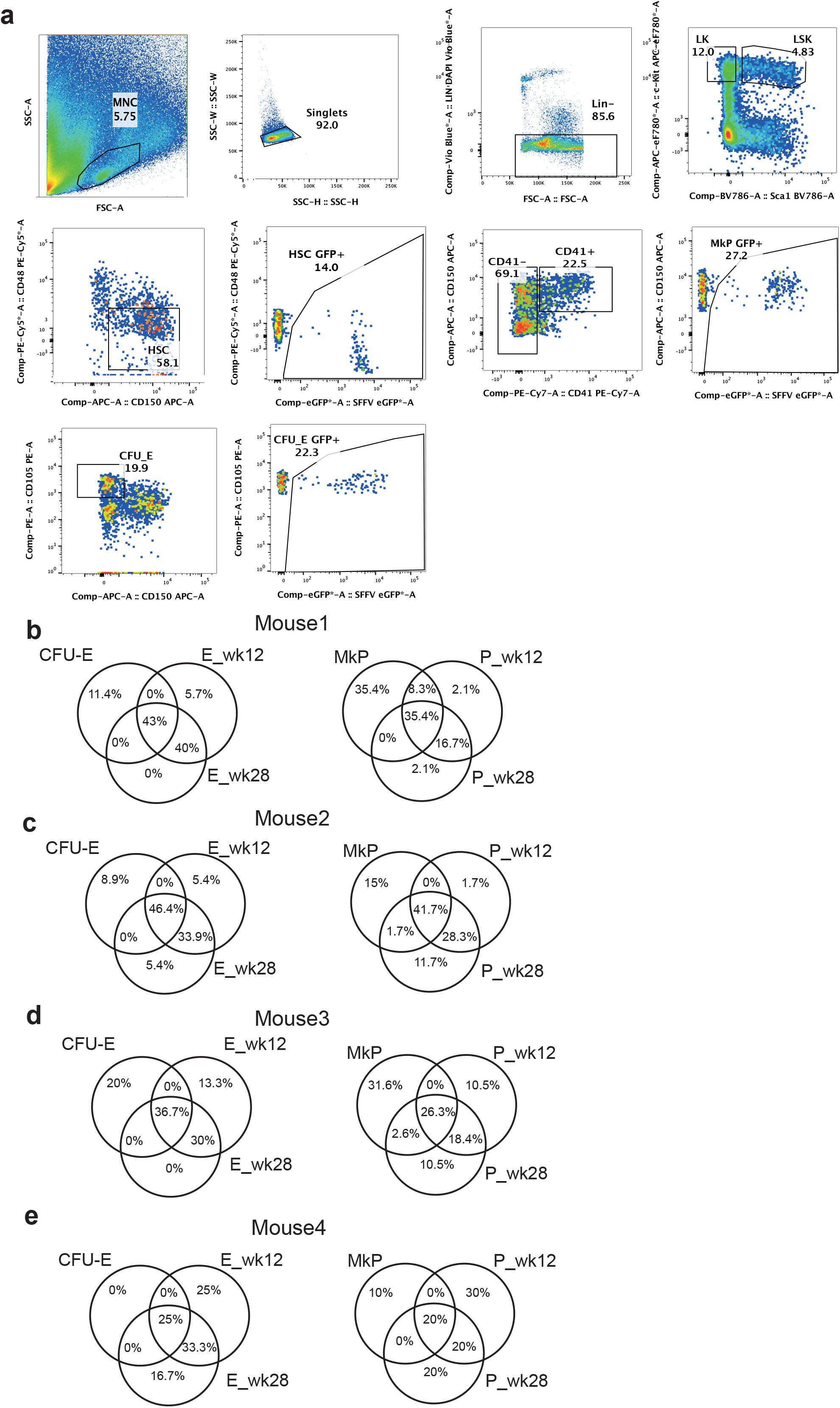
A. Gating strategy for FACS-purification of single BM populations: HSC, CFU-E and MkP, which were sorted as GFP^+^. B.-E. Venn diagrams representing the overlap between dominant barcodes (top 90% barcodes identified in all PB samples at 12 and 28-weeks post transplantation) detected in anucleate cells (platelets or erythroid cells) and their BM progenitors (MkP or CFU-E) in mouse 1-4

**Supplementary Figure 4.**
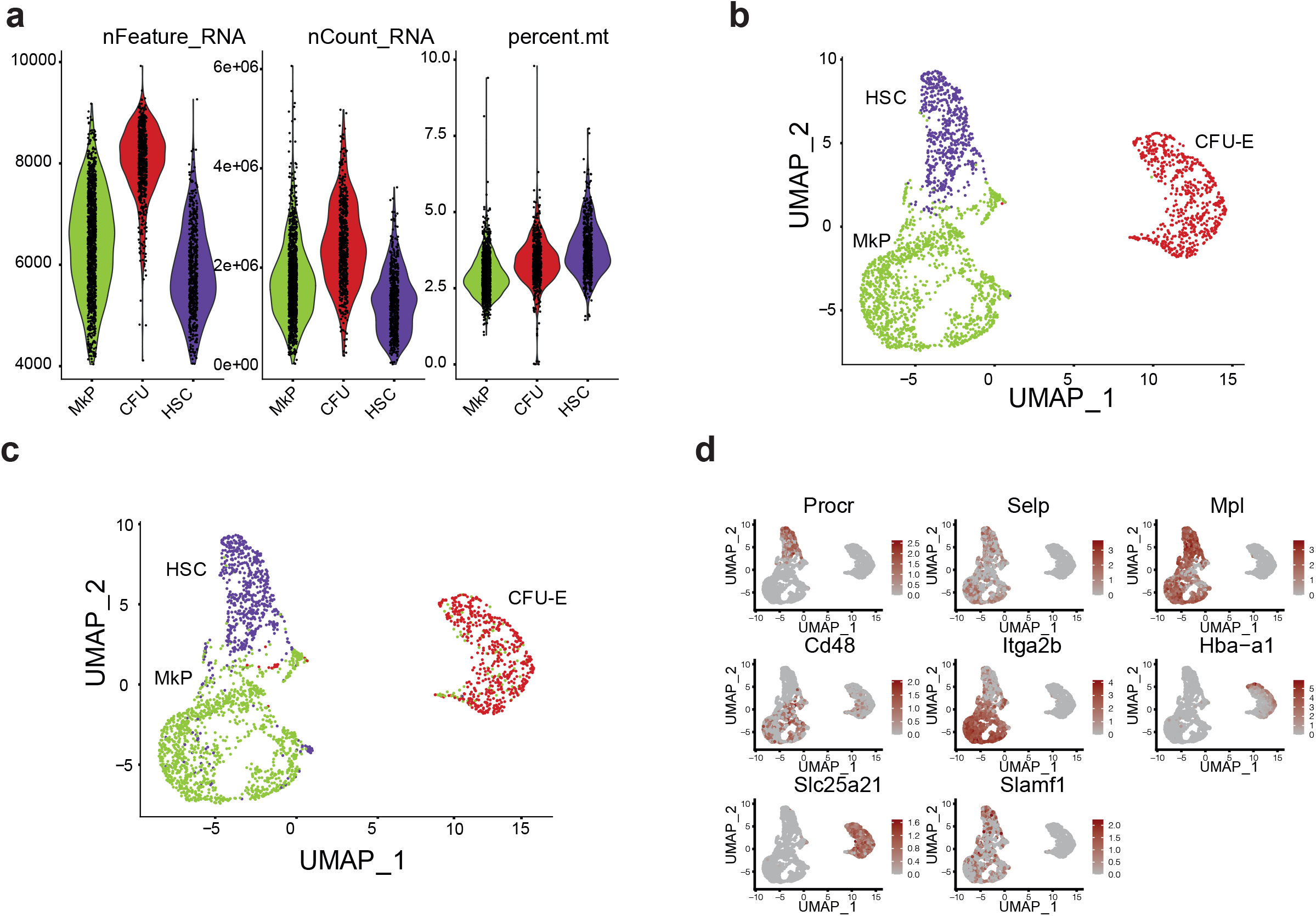
A. Violin plots showing the distribution of read count per cell, number of genes per single cell and fraction of mitochondrial genes in single cells within 3 analysed BM populations B. UMAP plot representing computationally assigned clusters of cells based on their transcriptomes (Seurat clusters) C. UMAP plot representing 3 clusters of cells split by their FACS phenotype D. UMAP plots representing the expression of marker genes for HSC (Procr, Slamf1, CD48, Mpl, Selp), MkP (Mpl, Slamf1, Selp) and CFU-E (Hba-a1, Slc25a21)

**Supplementary Figure 5.**
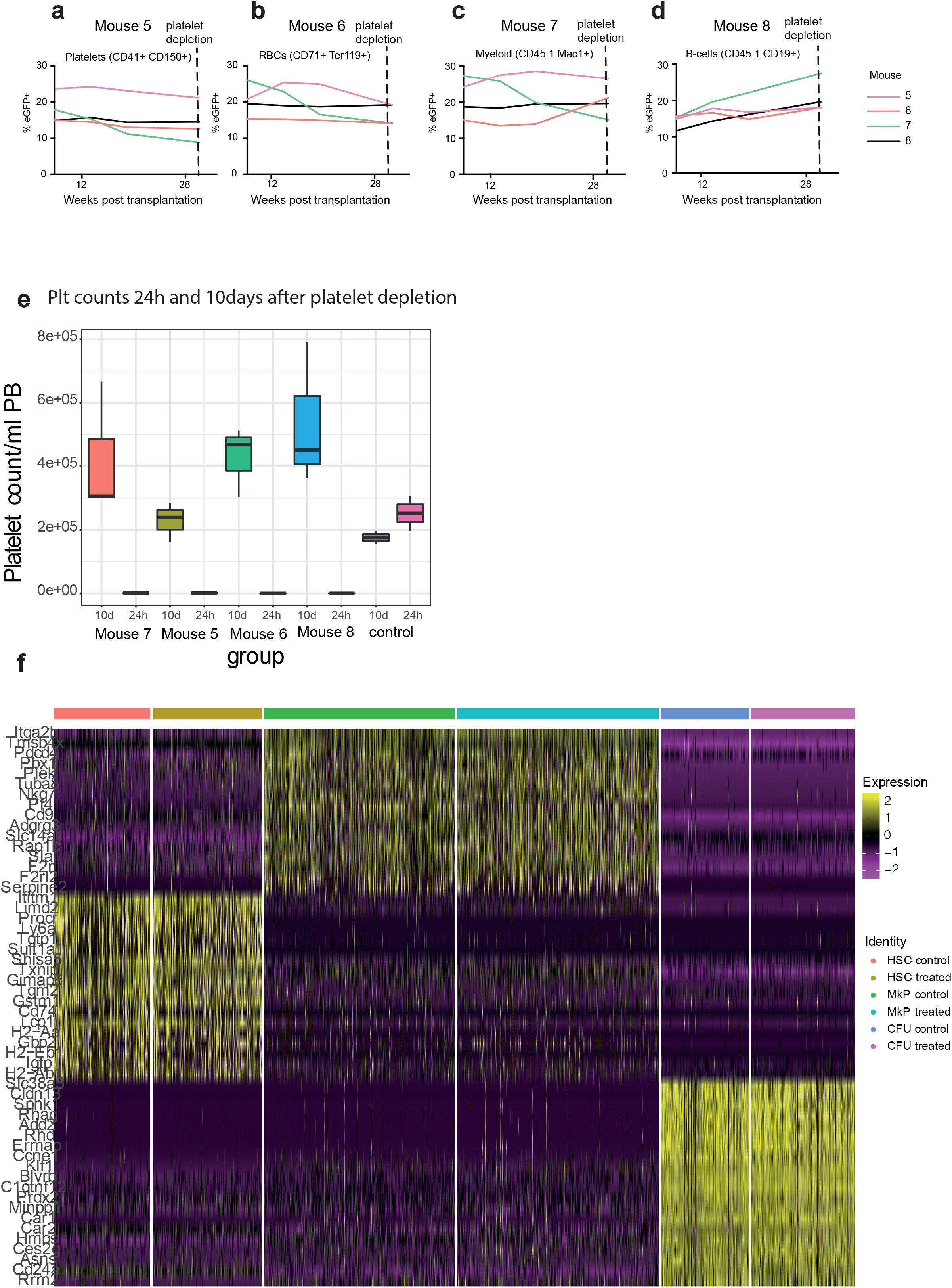
A. -D. Chimerism level in mature PB populations in mouse 5-8 E. Platelet count 24-hours and 10 days after platelet depletion in mouse 5-8 F. Expression heat map for BM cells (read count>50 000, mitochondrial gene content<10%) for the top 20 genes enriched in each cluster are displayed, showing gene expression on a log_2_ scale from yellow to purple, cell populations are split by experimental group (control/treated).

**Supplementary Table 1.**
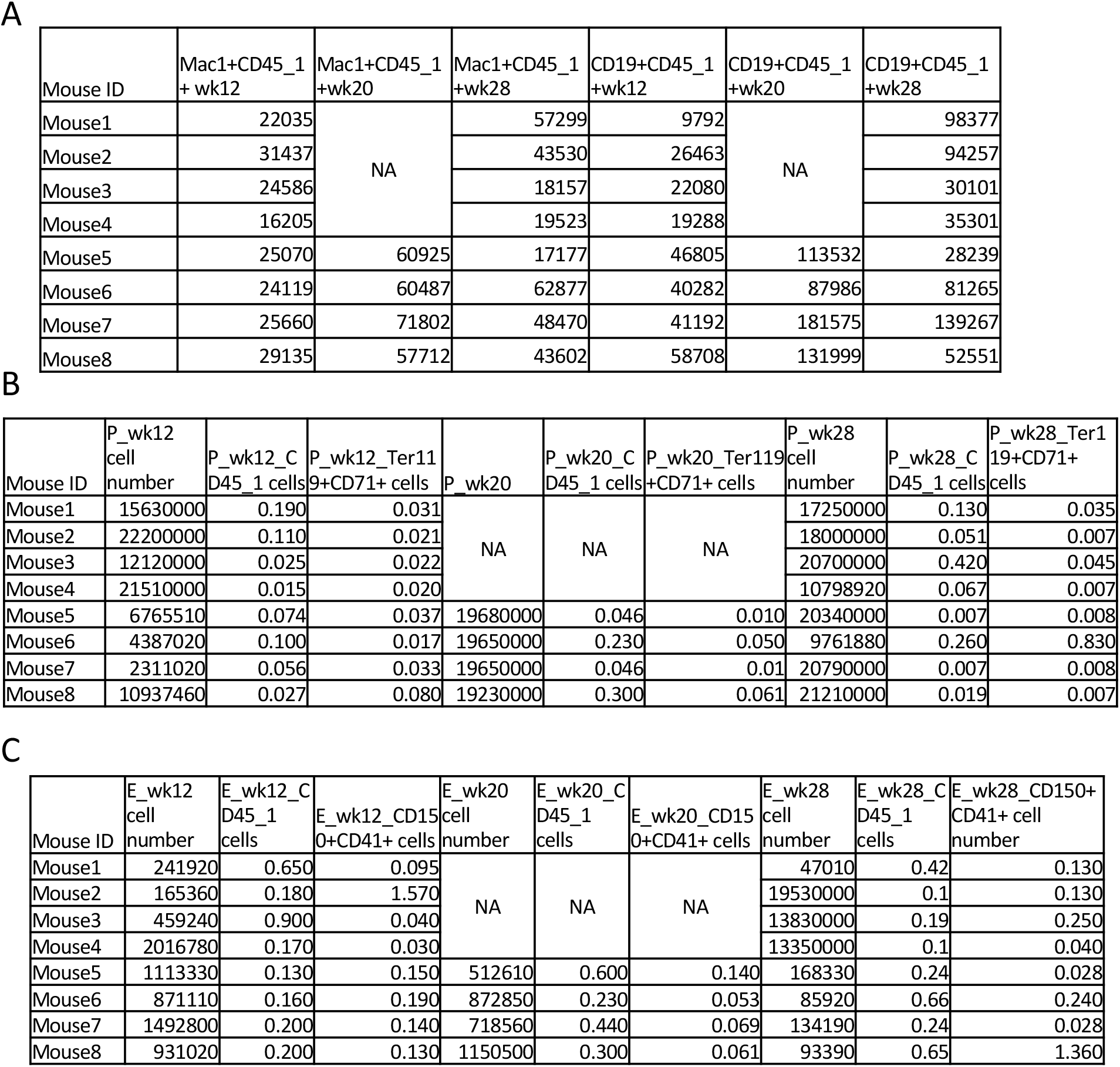
A-C Cell numbers for nucleated and anucleated cell populations used for barcode retrieval. The purity for auncleate cells (% of CD45.1+cells and Ter119+ for P-platelets, %CD45.1+cells and CD150+CD41+ cells for E-red blood cells) was evaluated by FACS.

**Supplementary Table 2.** Calculation of the smallest clone contributing to PB lineages: Counts for dominant barcodes (top 90%) were normalised per 1000 to account for the total read count/sample, next the size of each clone was recalculated to account for the actual eGFP chimerism level in different PB lineages at various time points (Suppl. Fig. 3 E-H, Suppl. Fig 6 A-D), consecutively the median was calculated from both technical replicates in each lineage at a given time point.

| biggest_<br>clone | clone type | contribution to<br>all lineages at<br>week 28 | smallest_<br>clone_P | clone<br>type | size | smallest_c<br>clone_E | clone<br>type | size | smallest_c<br>clone_M | clone<br>type | size | smallest_c<br>clone_B | clone<br>type | size |
| --- | --- | --- | --- | --- | --- | --- | --- | --- | --- | --- | --- | --- | --- | --- |
| mouse1 | PEMB | 33.05 | mouse1 | PE | 0.083 | mouse1 | EB | 0.026 | mouse1 | PEM | 0.13 | mouse1 | EB | 0.026 |
| mouse2 | PEMB | 10.6 | mouse2 | PE | 0.1 | mouse2 | PE | 0.1 | mouse2 | PMB | 0.3 | mouse2 | B | 0.08 |
| mouse3 | PEMB | 16.5 | mouse3 | PEM | 0.66 | mouse3 | EB | 0.011 | mouse3 | MEB | 0.5 | mouse3 | B | 0.77 |
| mouse4 | PEMB | 72 | mouse4 | PEMB | 1.77 | mouse4 | EB | 0.05 | mouse4 | MEB | 0.86 | mouse4 | MEB | 0.86 |
| mouse5 | MEB | 8.9 | mouse5 | PEMB | 0.17 | mouse5 | EB | 0.11 | mouse5 | MEB | 0.007 | mouse5 | EB | 0.011 |
| mouse6 | PEB | 14.3 | mouse6 | P | 0.0061 | mouse6 | EB | 0.0081 | mouse6 | PEM | 0.016 | mouse6 | MEB | 0.13 |
| mouse7 | MEB | 15.9 | mouse7 | P | 0.1 | mouse7 | MEB | 0.057 | mouse7 | MEB | 0.057 | mouse7 | MEB | 0.057 |
| mouse8 | PEMB | 22 | mouse8 | P | 0.079 | mouse8 | EB | 0.016 | mouse8 | PEM | 0.12 | mouse8 | EB | 0.016 |

**Supplementary Table 3.** Numbers of eGFP+ HSC, MkP and CFU-E cells recovered per mouse using SmartSeq2 protocol, which had over 50 000 reads/cell and less than 10% mitochondrial gene content, barcode read count>3.

|  | control |  |  |  | platelet depleted |  |  |  |
| --- | --- | --- | --- | --- | --- | --- | --- | --- |
|  | mouse1 | mouse2 | mouse3 | mouse4 | mouse1 | mouse2 | mouse3 | mouse4 |
| MkP | 168 | 166 | 168 | 166 | 162 | 174 | 156 | 178 |
| CFU | 68 | 70 | 80 | 69 | 97 | 86 | 78 | 87 |
| HSC | 76 | 74 | 60 | 58 | 82 | 76 | 80 | 90 |

**Supplementary Table 4.**
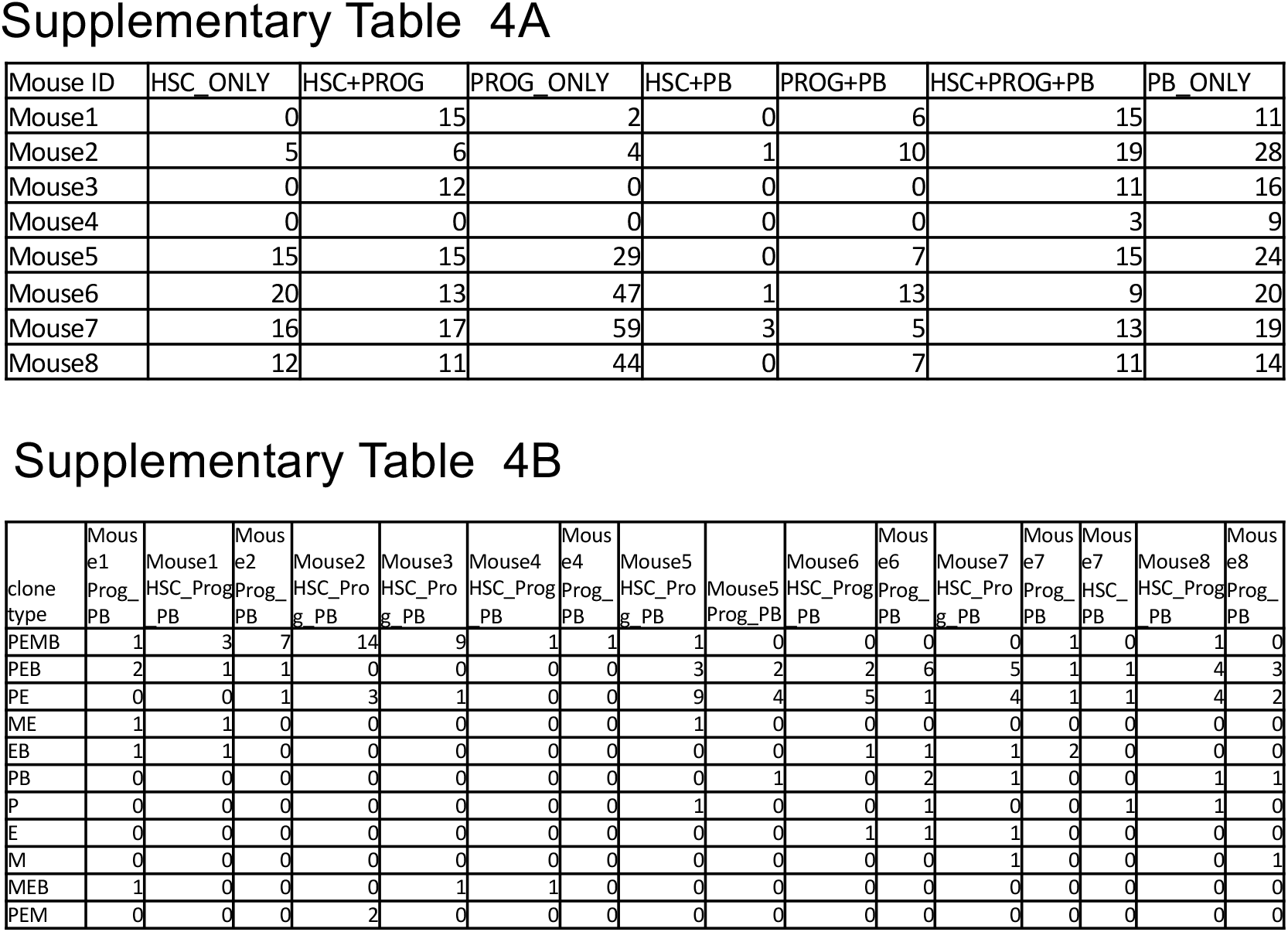
A. Numbers of clones found in HSC (HSC only), Progenitors (PROG only), HSC and Progenitors (HSC+PROG), HSC and PB (HSC and peripheral blood), Progenitors and PB (PROG+PB) or in all 3 HSC+Progenitors+PB (HSC+PROG+PB) to consider the clone as present in PB it had to contribute >0.1% to a lineage at 2 time points, otherwise clone would have been excluded from the analysis B. Types of clones found in HSC and PB, Progenitors and PB (PROG+PB) or in all 3 HSC+Progenitors+PB (HSC+PROG+PB) based on their output in PB, clone type name as in Fig.1K

**Supplementary Table 5.**
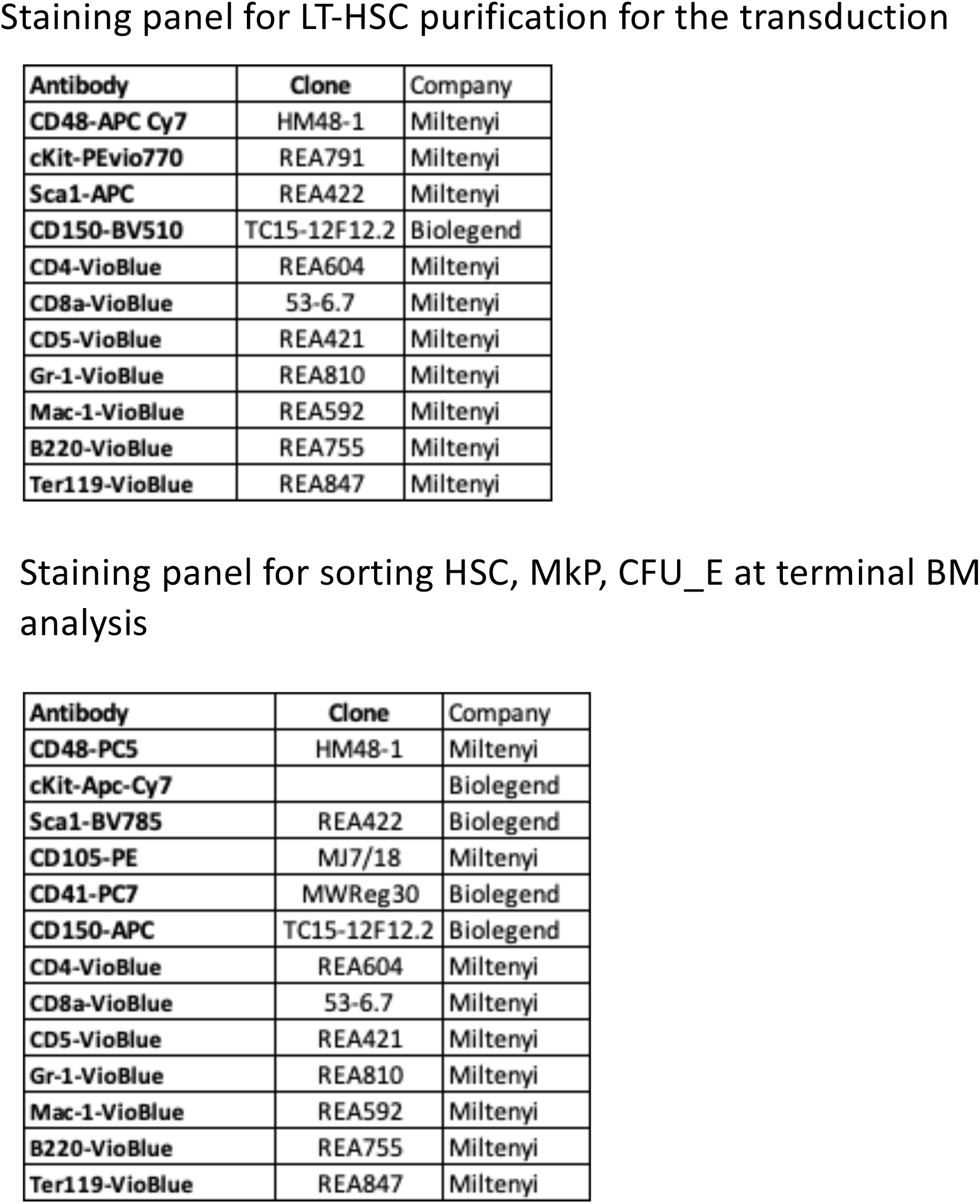
Staining panels for BM including antibody clone and conjugate.

**Supplementary Table 6.** Staining panels to FACS purify nucleated PB cells or evaluate the purity of anucleated cell populations.

Staining panel for sorting Mac1<sup>+</sup> and CD19<sup>+</sup> cells from PB
|  | Antibody | Clone | Manufacturer |
| --- | --- | --- | --- |
| MNC panel | Mac1-APC | m1/70 | Biolegend |
|  | CD150-BV785 | TC15-12F12.2 | Biolegend |
|  | CD4-APCe780 | RM4-5 | ThermoFisher |
|  | CD8-APCe780 | 53-6.7 | ThermoFisher |
|  | CD19-PC5 | eBioD3 | ThermoFisher |
|  | CD41-PC7 | MWReg30 | Biolegend |
|  | PE CD45.1 | A20 | Biolegend |
|  | PB CD45.2 | 109 | Biolegend |

Staining panel for evaluating the purity of platelets
|  | Antibody | Clone | Manufacturer |
| --- | --- | --- | --- |
| Platelet purity | Ter119-APC | Ter119 | Biolegend |
|  | CD41-PC7 | MWReg30 | Biolegend |
|  | CD45.1-PE | A20 | Biolegend |

Staining panel for evaluating the purity of erythroid cells
|  | Antibody | Clone | Manufacturer |
| --- | --- | --- | --- |
| RBC purity | Ter119-APC | Ter119 | Biolegend |
|  | CD41-PC7 | MWReg30 | Biolegend |
|  | CD150-BV785 | B217464 | Biolegend |
|  | CD71-BV421 | C2 | Biolegend |
|  | CD45.1-PE | A20 | Biolegend |

## Author contribution

EEW conceived the study, designed and performed experiments, analysed data and wrote the manuscript. VU performed the analysis of barcode data. JM, DWC, LK, AL, AS, FG, CH performed experiments. PSW designed experiments, revised the manuscript. MEB provided the barcoding library and revised the manuscript. GE performed the initial barcode extraction from fastq files. WH developed the pipeline for barcode retrieval in single cells, analyzed data and revised the manuscript. LVB provided the barcoding library, analyzed data and revised the manuscript. CN, KB designed experiments, revised the manuscript. SEJ conceptualized the study, designed experiments, analysed data and revised the manuscript. SAR designed and performed experiments, analysed data and revised the manuscript. ICM designed experiments, analysed data and wrote the manuscript.

### Funding

EEW is supported by a Sir Henry Welcome Postdoctoral Fellowship (213731/Z/18/Z), an Earlham Institute BBSRC Flexible Talent Mobility Fellowship (BB/R50659X/1) and previously by the Swedish Childhood Cancer Foundation Postdoctoral Fellowship at Karolinska Institutet, Sweden. Work at EI was supported by the Biotechnology and Biological Sciences Research Council (BBSRC), part of UK Research and Innovation, through the Core Capability Grant BB/CCG1720/1, BS/E/T/000PR9818 (ICM, WH, GE, VU), and at the Earlham Institute (ICM, WH, GE, VU) and BBS/E/T/000PR9816 (ICM, GE). AS was supported by the BBSRC funded Norwich Research Park Biosciences Doctoral Training Partnership grant BB/M011216/1. ICM is additionally supported by BBSRC New Investigator Grant BB/P022073/1. WH was additionally supported by a UK Medical Research Council [MR/P026028/1] award. Next-generation sequencing was delivered via the BBSRC National Capability in Genomics and Single Cell Analysis (BBS/E/T/000PR9816). JM was supported by BigC Cancer Charity. CH is funded by Wellcome Trust Clinical Research Fellowship. AML was funded by the Swedish Childhood Cancer Foundation Postdoctoral Fellowship. This work was supported by the The Swedish Research Council (538-2013-8995 to S.E.W.J.) and The UK Medical Research Council (MC_UU_12009/5 to S.E.W.J.).

### Data and materials availability

All data are available in the main text or the supplementary materials.

The GEO submission number associated with raw data presented in this manuscript is: GSE188268

## Supplementary Materials Methods

### Mice

B6.SJL-Ptprca^Pep3b/BoyJ^ (CD45.1) mice were purchased from The Jackson Laboratory (Bar Harbor, ME). C57BL/6J mice (CD45.2), B6.129S-Cybb^tm1Din/^J mice were purchased from Charles River Laboratories. Animals were housed in a specific pathogen-free facility. All animal work in this study was carried out in accordance with regulations set by the United Kingdom Home Office and the Animal Scientific Procedures Act of 1986. Donor mice were 8 to 12 weeks old, recipient animals were 6-12 weeks old, and both genders were used for experiments.

### Cell isolation and preparation

BM isolation was prepared by isolating the tibia, femur, pelvis, spine and sternum of each mouse. Bones were crushed in FACS media (phosphate buffer solution (PBS, ThermoFisher, MA, USA) supplemented with 2mM EDTA (Sigma Aldrich, Darmstadt, Germany) and 5% fetal bovine serum (Invitrogen, Darmstadt, Germany)). Stem and progenitor cells were enriched from BM using CD117 (cKit) microbeads (Miltenyi Biotech, Bergisch Gladbach, Germany) prior to staining and sorting.

### Cell sorting

Antibody mixtures were prepared in 1 x FACS media and incubated with BM cells for 15-30 min at 4 °C. For flow cytometric cell sorting of BM cell populations, cells were resuspended in antibody mix, and cells were sorted directly into StemSpan (Stem Cell Technologies, Canada) with penicillin and streptomycin (GIBCO, MA, USA), hTPO, mSCF (Peprotech, MA, USA) or into Smart-seq2 lysis buffer. Cell sorting was performed on a BD FACS Melody (BD Bioscience, CA, USA). See Supplementary Fig. 2 for specific gating strategies, staining panel Supplementary Table 5.

### Virus production

HEK293T/17 cells (ATCC, Virginia, USA) used for barcoding library virus product were cultured in DMEM (Thermo Fisher Scientific, Massachusetts, USA) with 10% FSC (Thermo Fisher Scientific), penicillin and streptomycin (GIBCO, MA, USA) and incubated in 37°C, in 5% CO_2_, with R95% humidity.

To generate the pEGZ2 lentiviral barcoding library[18] HEK293T cells (ATCC, Virginia, USA) were transfected with the pGIPZ-based library, pCMV-Δ8.9 and Vsv-g plasmids using Lipofectamine 2000 (Thermofisher, MA, USA) in Opti-MEM (Thermofisher, MA, USA). 24-hours post transfection medium was changed to DMEM (Thermo Fisher Scientific, MA, USA) with 10% FSC (GIBCO, MA, USA), penicillin and streptomycin (GIBCO, MA, USA). Harvests were collected 48 and 72h post transfection, and concentrated by centrifugation (8 h at 6000 g, 4°C). Cells were transduced with a barcoding library at an MOI of 50, defined as the titre on BaF3[37] cells divided by the number of HSCs. This generated an HSC infection rate of ca. 45%.

### Transduction and transplantation

LT-HSC were sorted into StemSpan medium (StemCell Technologies, Vancouver, Canada) supplemented with 10ng/ml mouse recombinant SCF, 100ng/ml human TPO (both Peprotech, MA, USA) and penicillin/streptomycin (ThermoFisher, MA, USA). Cells were spun down and resuspended in the viral supernatant[18] supplemented with the same concentrations of mSCF, hTPO, antibiotics and polybrene (5ug/ml, Sigma Aldrich, Darmstadt, Germany) was added. Cells were harvested 15-20 hours later, extensively washed, counted and resuspended in transplantation media (PBS + 5% FBS). Cells were immediately injected intravenously into mice (8-10 weeks old) preconditioned with busulfan 25 mg/kg/day for 3 days. The transduction efficiency was 47% (±17%) in both experiments.

For evaluation of eGFP silencing LSK cells were FACS-sorted (BD FACS Aria Fusion, CA, USA) from VavCre x Rosa26tdTomato 8-10 week old donors and transduced with barcoded viruses. 30-hours after transduction eGFP+ viable cells were FACS-sorted (BD Fusion, CA, USA) and approximately 2,500 eGFP+ tdToamto+ cells were transplanted into 8-12 week old CD 45.1 lethally irradiated (960rad given as a split dose) recipient mice along with 200 000 CD45.1+ whole BM support cells.

### Peripheral blood cell isolation

Mice were bled every 6-8 weeks. Blood was collected to EDTA coated tubes (Sarstedt, Numbrecht, Germany), shaken vigorously and kept at room temperature. Next, tubes were spun down at 200g for 8 minutes at room temperature with no acceleration or brake. Platelet rich plasma fraction was collected and used for platelet purification. The lower RBC fraction and the buffy coat were transferred to a new tube and mixed with 2% Dextran (Sigma Aldrich, Darmstadt, Germany) in 1:1 volumetric ratio with FACS media. A small fraction (2ul) of the lower fraction was resuspended in isolation media (PBS+0.1%BSA, 2mM EDTA) and used for the purification of erythrocytes. Cells with Dextran were incubated for 15 minutes at 37°C. Next, the upper phase was washed in FACS media and spun down (5 minutes, 500g, 4°C). The supernatant was aspirated and the cell pellet resuspended in ammonium chloride (Stem Cell Technologies, BC, Canada), incubated at room temperature, washed and stained with antibody mix in Supplementary Table 6.

Cells were stained for 15-30 minutes at 4°C, washed in FACS media and sorted on BD FACS Melody (BD, CA, USA). Sorted myeloid cells were defined as Mac1^+^ (CD19^-^CD4/8^-^ CD41^-^ CD150^-^ Mac1^+^ SSC^hi^ CD45-1^+^), CD19+ (Mac1^-^ CD4/8^-^ CD41^-^ CD150^-^ CD19^+^ CD45-1^+^).

Platelet rich plasma was resuspended in 100 μl of isolation media - PBS supplemented with 0.1%BSA, 2mM EDTA, 0.001M prostaglandin E2 (Millipore, Darmstadt, Germany) and 0.02U/ml apyrase (Merck, Darmstadt, Germany) and stained with biotin-anti-mouse Ter119 antibody (clone-TER-119) and biotin-anti-mouse CD45 antibody (clone 30-F11, both from Thermo Fisher, MA, USA) for 15 minutes. Cells were washed in isolation media, spun down (5 minutes, 22°C, 450g) and resuspended in 100 ul of isolation media with Biotin Binder Dynabeads (Thermo Fisher, MA, USA). Cells were incubated 15 minutes on the rotator at room temperature. Next, cells were placed on the magnet, the supernatant was transferred to a new tube. 3-5 ul of the supernatant was subjected for the purity test by FACS using the following antibodies on BD FACS Melody, Supplementary Table 6.

Erythrocyte suspension was incubated with anti-mouse biotin CD41 (clone MWReg.30) and anti-mouse-biotin CD45 (clone 30-F11, both Thermo Fisher, MA, USA) for 15 minutes. Next, cells were washed in the FACS media, supn down and resuspended in 100ul of FACS media with Dynabeads Biotin Binder (Thermo FIsher, MA, USA). Cells were incubated for 15 minutes on the rotator and placed on the magnet. The supernatant was transferred to the new tube, 2-3 ul of cells were used for the purity test using BD FACS Melody and indicated antibodies (Supplementary Table 3).

### Platelet depletion

Mice transplanted with barcoded LT-HSC were intravenously injected with anti CD42b antibody (R-300, Emfret, Mainz, Germany) at a dose 2μg/g body weight 28-weeks post transplantation. Blood samples were taken 24 hours post depletion to measure platelet count. A small blood sample was obtained by tail vein puncture with a 26G needle, blood was collected in EDTA-treated tubes (Sarstedt, Numbrecht, Germany), stained with CD41-PC7 and run on FACS Melody with Sphero Counting Beads (Spherotech, IL, USA). Animals were sacrificed 10 days later and peripheral blood and bone marrow cells were subjected for the clonal analysis.

### Single cell RNA sequencing-library construction

Single cells from the bone marrow (LT-HSC, MkP, CFU-E) were FACS sorted (BD FACS Melody) into 2 μl Smart-seq2 lysis buffer[26]. Plates were stored at -80 until further processing according to standard Smart-seq2 protocol[26]. Reverse transcription conditions: 42°C for 90 minutes, 10 cycles 50°C for 2 min, 42°C for 2 min, hold at 4°C; PCR amplification conditions: 98°C for 3 min, 21 cycles: 98°C for 20 sec, 67°C for 15 sec, 72°C for 6 min, final extension 72°C for 5 min, hold at 4°C. Amplified cDNA underwent bead clean-up in a 1:0.8X volumetric ratio of Ampure Beads (Beckman Coulter, FL, USA) and eluted into 25 μl of elution buffer (EB, Qiagen, MD, USA). Selected libraries were analysed on the Bioanalyzer using a High Sensitivity DNA chip (Perkin Elmer, MA, USA) to assess their quality. Next, cDNA was used for library construction using Nextera XT Kit (Illumina, CA, USA). Resulting libraries were pooled in batches of 384 and cleaned up using a 1:0.6 volumetric ratio of Ampure Beads (Beckman Coulter, FL, USA) and eluted into 25 μl of elution buffer (EB, Qiagen, MD, USA). Samples were barcoded during library preparation and 150bp paired-end reads were generated on a NovaSeq6000 (Illumina, CA, USA).

### Blood sample processing for barcode recovery

Blood cell populations were sorted into FACS media, spun down, resuspended in RLT Plus Buffer (Qiagen, MD, USA) and stored at -80°C. Lysates were slowly thawed on ice and incubated with streptavidin coated magnetic beads (DynaBeads, ThermoFisher, MA, USA) as previously described[20]. The original protocol has been modified-for barcoded transcript capturing on beads we have used a biotinylated barcode specific primer (Rev6 5’- CGTCTGGAACAATCAACCTCTGG-3’). Obtained cDNA was used for the barcode amplification using forward primer containing a 9 nucleotide tag for multiplexing (Fwd3 5’NNNNNNNNNCGGCATGGACGAGCTGTACAAG-3’) on the thermal cycler as follows: 95 °C for 3 min, then 30 cycles of 95 °C for 20 s, 63 °C for 15 s, 72 °C for 9 s and finally 72 °C for 5 min. Amplified cDNA was cleaned up using a double sided SPRI 1:1.2, collected the supernatant, transferred to a new tube and added 1:2.5X volumetric ratio of Ampure Beads (Beckman Coulter, FL, USA) and eluted into 25 μl of elution buffer (Buffer EB, Qiagen, MD, USA).

Libraries were used for PCR-free ligation of adapters (TruSeq, Illumina, CA, USA) following manufacturer protocol with the modification for the bead clean up ratio -used 1:2X volumetric ratio (Ampure XT beads, Beckman Coulter, FL, USA). Obtained libraries were evaluated on Bioanalyzer and sequenced on NovaSeq6000 (150bp, paired-end, Illumina, CA, USA).

## Barcode recovery

### Single cells

Obtained from the sequencer Fastq files were used for alignment to *Mus musculus* transcriptome version M38 using STAR aligner and count reads[38]. To obtain a single-cell expression matrix object we used a custom R script. Subsequent analysis was performed in R using Seurat version 4.0.0[39]. Cells showing gene counts lower than 50000 reads and a mitochondrial gene expression percentage higher than 10% were excluded from further analysis. Within Seurat, data were normalised using the NormalizeData function (using the LogNormalize method and scale.factor = 10000). The barcode sequence was recovered from scRNA-seq dat using BBduk which located the eGFP primer with Hamming distance=1 (+ or - 1 nucleotide), subtracted this sequence and extracted the barcode structure. Barcode sequence and count were summarised in a table. All barcodes with the read count below 3 were ignored. The probability of capturing 4 reads for the same barcode in 1 well purely by chance with the library consisting of 800 barcodes is (1/800)^4=2.44E-12, thus we considered it highly unlikely and applied in our pipeline. Detected barcode list is included in Supplementary Table 7.

### Bulk PB sample barcode recovery

Barcodes for the PB samples were retrieved using custom scripts in python, as described previously[21]. We have detected 1 noninformative barcode (sequence: 5’- AAGGGCAACCTGGTAACCGATCTATGACACGATGTGTGACGGC-3’), which was excluded from all further analysis.

To obtain the Shannon count we firstly calculated the Shannon index (H):

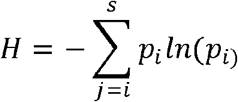

where *s* is the total number of observed barcodes in the sample, *p* is the proportion of reads belonging to the *i*-th barcode in the sample. Next, the Shannon index is converted into the Shannon count:

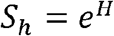

If all barcodes are equally distributed in the sample the Shannon count is equal to the number of all barcodes detected in the sample. However, when the barcode distribution is skewed and their contribution is unequal Shannon count is lower than the number of all barcodes, which makes Shannon count less sensitive to PCR noise[40]. The list of barcodes detected in individual animals is attached in Supplementary Table 8.

### Statistical analysis

Consistency in barcode patterns was expressed as Spearman correlation. Pearson correlation was applied as a measure of the distance between PB samples. Differences between groups were calculated using Student t-test for normally distributed variables.

## Figure legends

**Supplementary Figure 1**

A. Experimental set up for *in vitro* validation of quantitative RNA based clonal studies. The K562 cell line was used for the transduction, single eGFP+ cells were FACS-sorted and expanded to generate monoclonal calibration samples. Four clones were used to create samples, where cells carrying different barcodes were mixed in known ratios-spiked in clone contributing 0.1%, 1%, 5%, 10%, 20% or 50% of total mix.

B. Generalised linear model (GLM) correlation between the read count for barcoded transcript retrieved in calibration samples from cDNA or gDNA.

C. Experimental setup to test the silencing of eGFP in VavCre x Rosa26tdTomato LSK cells

D.-H. eGFP+ chimerism changes in tdTomato+ anucleate cells and CD45.2+ (donor derived) nucleated cells. To calculate the silencing in specific lineage we subtracted eGFP^+^ cells from CD45.2^+^ cells (nucleated cells-CD19^+^, Mac1^+^ or CD4/8^+^cells) or from tdTomato^+^ cells (all donor derived anucleate cells), shown are results from a single experiment in 3 recipient mice

I. Gating strategy to purify HSC (LSK CD150^+^48^-^) for transduction and transplantation experiments (4-6 mice used as donors in each experiment)

J.-M. Contribution of transduced, eGFP^+^ HSC into 4 mature PB populations during the course of the experiment, plots are showing the chimerism level in mice 1-4 (corresponds to clonal composition in mice depicted in Fig.1 D-G)

N.-R. Linear regression between 2 technical replicates for cDNA retrieved from 4 mature PB populations (P-platelets, E-erythroid cells, M-myeloid cells, B-B-cells) to assess the reproducibility of barcode recovery (displayed are normalised barcode read count in 2 technical replicates in the same animal)

**Supplementary Figure 2**

A. Gating strategy for PB cell population FACS-purification. Displayed are gates within the MNC/singlet/viable cell population. FACS definitions for purified populations: Mac1^+^ (CD19^-^CD4/8^-^CD150^-^CD41^-^CD45.1^+^), CD19^+^ (Mac1^-^CD4/8^-^CD19^+^CD150^-^CD41^-^CD45.1^+^)

B. Purity of sorted Mac1^+^ cells evaluated by FACS

C. Purity of CD19^+^ cells evaluated by FACS

**Supplementary Figure 3**

A. -D Stacked bar plots representing clonal composition in PB populations (P-platelets, M-myeloid cells, E-erythroid cells, B-B cells) at 12 and 28 weeks in 4 analysed control animals, on top depicted is the Shannon count.

E.-H. Contribution of clones (top 90% of all barcodes detected in PB) corrected for the chimerism level in analysed PB lineages (Suppl. Fig 1 J-M)

I. - K. MDS with Pearson correlation between PB cell types (P-platelets, E-erythroid cells, M-myeloid cells, B-B cells) at week 12 and 28 in mouse 1, 3 and 4

**Supplementary Figure 4**

A. Gating strategy for FACS-purification of single BM populations: HSC, CFU-E and MkP, which were sorted as GFP^+^.

B. -E. Venn diagrams representing the overlap between dominant barcodes (top 90% barcodes identified in all PB samples at 12 and 28-weeks post transplantation) detected in anucleate cells (platelets or erythroid cells) and their BM progenitors (MkP or CFU-E) in mouse 1-4

**Supplementary Figure 5**

A. Violin plots showing the distribution of read count per cell, number of genes per single cell and fraction of mitochondrial genes in single cells within 3 analysed BM populations

B. UMAP plot representing computationally assigned clusters of cells based on their transcriptomes (Seurat clusters)

C. UMAP plot representing 3 clusters of cells split by their FACS phenotype

D. UMAP plots representing the expression of marker genes for HSC (Procr, Slamf1, CD48, Mpl, Selp), MkP (Mpl, Slamf1, Selp) and CFU-E (Hba-a1, Slc25a21)

E. -H. Chimerism level in mature PB populations in mouse 5-8

**Supplementary Figure 6**

A. -D. Clonal contribution of barcoded cells to 4 mature PB lineages (P-platelets, E- erythroid cells, M-myeloid cells, B-B cells) corrected for the chimerism in these lineages (Suppl. Fig. 5 E-H) in platelet depleted animals (mouse 5-8), timepoint after platelet depletion labelled as #

E. Platelet count 24-hours and 10 days after platelet depletion in mouse 5-8

**Supplementary Figure 7** A. -C. MDS with Pearson correlation between PB (P-platelets, E-erythroid cells, M-myeloid cells, B-B cells) clone abundance for mouse 5, 7 and 8 after platelet depletion showing longest distance between myeloid cells at week 28+10days, # indicate the time point after platelet depletion

D. - G. Stacked bar plots representing the barcode composition of PB cell populations (P-platelets, E-erythroid cells, M-myeloid cells, B-B cells) in platelet depleted mice, on top depicted is the Shannon count, # indicates the time point after platelet depletion

**Supplementary Table 1** A-C Cell numbers for nucleated and anucleated cell populations used for barcode retrieval. The purity for auncleate cells (% of CD45.1+cells and Ter119+ for P-platelets, %CD45.1+cells and CD150+CD41+ cells for E-red blood cells) was evaluated by FACS.

**Supplementary Table 2** Calculation of the smallest clone contributing to PB lineages: Counts for dominant barcodes (top 90%) were normalised per 1000 to account for the total read count/sample, next the size of each clone was recalculated to account for the actual eGFP chimerism level in different PB lineages at various time points (Suppl. Fig. 3 E-H, Suppl. Fig 6 A-D), consecutively the median was calculated from both technical replicates in each lineage at a given time point.

**Supplementary Table 3** Numbers of eGFP+ HSC, MkP and CFU-E cells recovered per mouse using SmartSeq2 protocol, which had over 50 000 reads/cell and less than 10% mitochondrial gene content, barcode read count>3.

**Supplementary Table 4**A. Numbers of clones found in HSC (HSC only), Progenitors (PROG only), HSC and Progenitors (HSC+PROG), HSC and PB (HSC and peripheral blood), Progenitors and PB (PROG+PB) or in all 3 HSC+Progenitors+PB (HSC+PROG+PB) to consider the clone as present in PB it had to contribute >0.1% to a lineage at 2 time points, otherwise clone would have been excluded from the analysis

B. Types of clones found in HSC and PB, Progenitors and PB (PROG+PB) or in all 3 HSC+Progenitors+PB (HSC+PROG+PB) based on their output in PB, clone type name as in Fig.1K

**Supplementary Table 5** Staining panels for BM including antibody clone and conjugate.

**Supplementary Table 6** Staining panels to FACS purify nucleated PB cells or evaluate the purity of anucleated cell populations.

**Supplementary Table 7** Summary of barcodes found in the BM in single cells.

**Supplementary Table 8** Summary table of barcodes detected in PB populations.

